# F_420_ reduction as a cellular driver for anaerobic ethanotrophy

**DOI:** 10.1101/2024.01.23.576903

**Authors:** Olivier N Lemaire, Gunter Wegener, Tristan Wagner

## Abstract

The anaerobic ethane oxidation performed by seafloor archaea and sulfate-reducing partner bacteria involves largely uncharted biochemistry. This study deciphers the molecular basis of the CO_2_-generating steps by characterizing the native archaeal enzymes isolated from a thermophilic enrichment culture. While other microorganisms couple these steps to ferredoxin reduction, we found that the CO-dehydrogenase and the formylmethanofuran-dehydrogenase are bound to an F_420_-reductase module. The crystal structures of these multi-metalloenzyme complexes revealed a [4Fe-4S]-cluster networks electronic bridges coupling C1-oxidation to F_420_-reduction. Accordingly, both systems exhibit robust F_420_-reductase activities, which are not detected in methanogenic or methanotrophic relative organisms. We speculate that the whole catabolism of these archaea is reoriented towards F_420_-reduction, which facilitates the electron transfer to the sulfate-reducing partner, therefore representing the driving force of ethanotrophy.

## Introduction

Alkanes are the most reduced carbon compounds available in nature that can serve as cellular energy sources for numerous microorganisms in oxic and anoxic environments (1-3). In marine cold seeps and hydrothermal vents, anaerobic alkane oxidation prevents the release of alkanes to the oceans and sustains chemoautotrophic microorganisms through sulfide generation (4-6). Only two archaeal species belonging to the *Methanosarcinales* order can catalyze the complete anaerobic oxidation of ethane to CO_2_ (7-9). The electrons released during the oxidation process are transferred to sulfate-reducing bacteria. Ethane is initially activated as an ethyl-thiol adduct on the coenzyme M (CoM) via the ethyl-CoM reductase (ECR) (10). It has been suggested that the generated ethyl-CoM is further processed to acetyl-CoA based on the acquired knowledge of methanogens belonging to the same order, together supported by transcriptomics and proteomics data (7, 8). Based on the accepted metabolic model, the acetyl-CoA decarbonylase/synthase complex (ACDS) would transform the acetyl-CoA to generate CO_2_ concomitantly with a methyl-group branched on a tetrahydromethanopterin carrier (CH_3_-H_4_MPT, Fig. 1A). The methyl group would be oxidized through the reverse methanogenesis pathway to be ultimately released as CO_2_ by the formylmethanofuran dehydrogenase complex (Fmd/Fwd for molybdenum/tungsten-dependent enzymes, Fig. 1A) (11-13). Both CO_2_-releasing steps would be coupled to ferredoxin reduction, which is employed for energy conservation in methanogens (14). Ethanotrophs do not contain any known membranous systems that would allow energy conservation from ferredoxin oxidation, questioning the dogma of the CO_2_-releasing step coupled to ferredoxin in these organisms. To solve this metabolic puzzle, we focused our interest on the highly sophisticated multi-enzymatic ACDS and Fwd/Fmd complexes by isolating both enzymes directly from a microbial enrichment (15-22).

**Figure 1.**
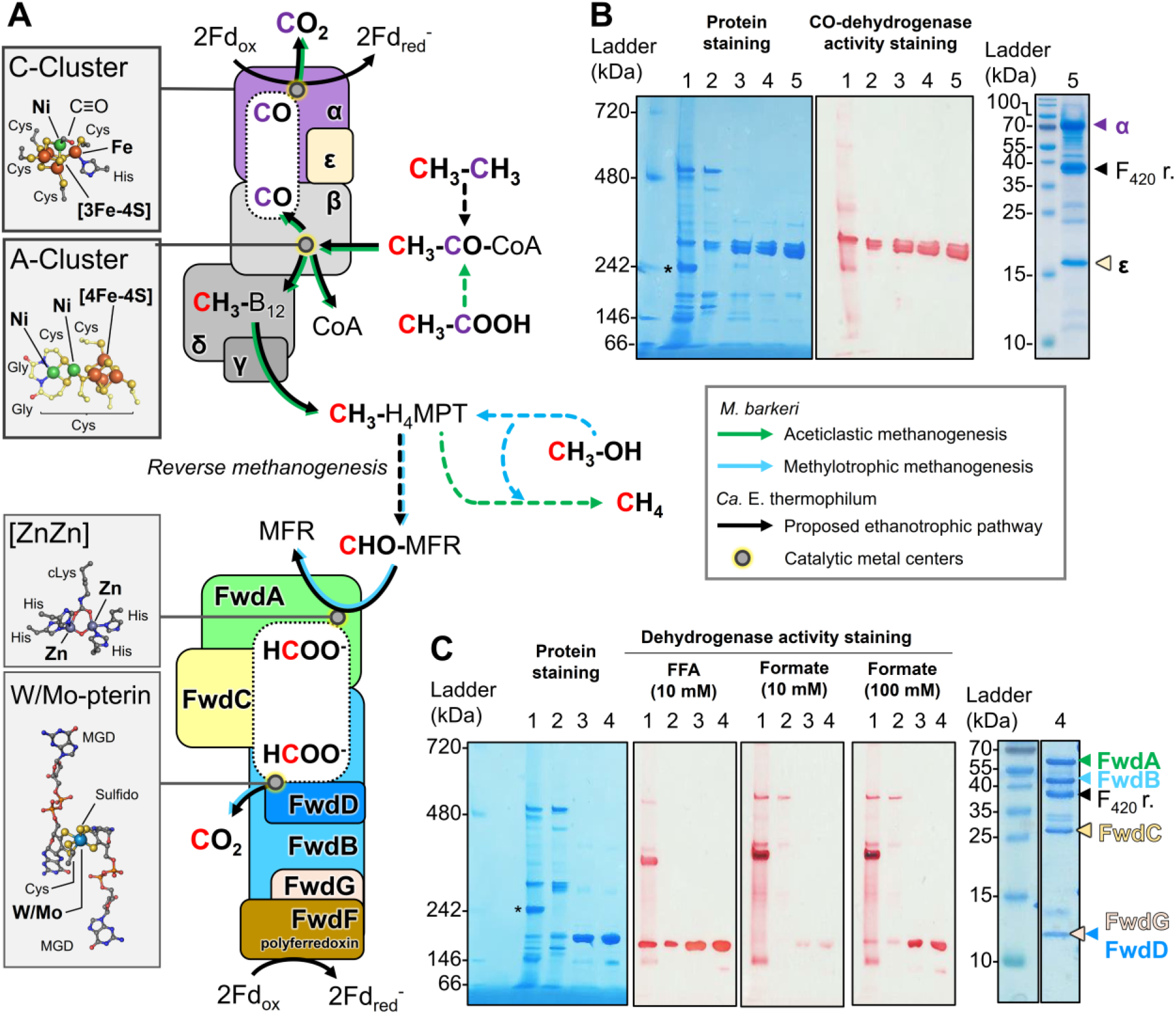
Proposed catabolic main pathway of *Ca*. E. thermophilum, and native purification of the CODH and Fwd/Fmd complexes. **A**. The pathway is proposed based on studies described in *Methanosarcinales* (7, 8, 10). The assembly of the ACDS (top) and Fwd (bottom) complexes are drawn in compliance with previous studies (16, 18, 32). Arrows are colored according to the corresponding metabolism and dashed lines indicate multi-step transformations. Dotted lines illustrate internal channeling systems in which the substrates diffuse. Metallocofactor structures displayed in the insets are derived from the deposited PDB models 1RU3 (acetyl-CoA synthase from *Carboxydothermus hydrogenoformans* (33)), 3CF4 (carbonylated ACDS α_2_ε_2_ subcomplex from *M. barkeri* (16)) and 5T5M (Fwd complex from *M. wolfei* (18)) with metals in bold. cLys stands for modified carboxylysine. **B**. Purification steps of the CODH subcomplex on native PAGE (left). 1, soluble extract; 2, Anion exchange chromatography; 3, 4, hydrophobic exchange chromatography, and 5, size exclusion chromatography. **C**. Purification steps of the Fwd/Fmd complex on native PAGE (left). 1, soluble extract; 2, anion exchange chromatography; 3, hydrophobic exchange chromatography, and 4, size exclusion chromatography. **B** and **C**, a denaturating (right) PAGE of the enriched fractions. Subunits are indicated by their respective accession numbers. **B** and **C**, an asterisk marks the band corresponding to the Ethyl-CoM reductase (10) on the native electrophoresis profile. F_420_ r. stands for the additional F_420_-reductase subunit.

*Candidatus* Ethanoperedens thermophilum, the model ethanotrophic organism sampled from the gas-rich Guaymas Basin hydrothermal vent, was reported to be the fastest-growing anaerobic alkane oxidizer reported so far and represents around 40% of the microbial population in our enrichment culture (8). Purifying enzymes from such a heterogeneous microbial mixture would represent a challenge if these complexes were not abundant. However, published transcriptomic data confirmed that genes coding for the subunits of the ACDS and Fwd/Fmd complexes are among the 250 most expressed genes in the culture conditions. Their natural abundance in the cell extract was attested by the final yields obtained during their purification and observation on native PAGE (PolyAcrylamide Gel Electrophoresis, Fig. 1B and C, Table S1). The ACDS was followed by measuring the CO-oxidation activity carried by the CO-dehydrogenase subunit (CODH, α subunit) and Fwd/Fmd complex by the oxidation of the surrogate furfurylformamide instead of formylmethanofuran (CHO-MFR) (23). Fwd/Fmd activity was also tested against formate since it has been previously described that these enzymes exhibit a formate dehydrogenase activity, albeit at low rates and high formate concentration (Fig. 1C) (24). The artificial electron acceptor methyl-viologen was used as an electron acceptor for both complexes. The CO-oxidase and furfurylformamide oxidase activities were anaerobically enriched from the microbial enrichment through a five and four-step purification protocol, respectively (Table S1). The molecular weights of both purified complexes were estimated based on native PAGE and size exclusion chromatography (Fig. 1B and C, Fig. S1). In *Methanosarcina* species, the CODH is part of the 2.4 MDa ACDS composed of five subunits (α_8_β_8_γ_8_δ_8_ε_8_ stoichiometry) (15, 16), or a subcomplex α_2_ε_2_ of 215 kDa for which none would fit with the experimentally determined molecular weight (around 310 kDa) for the CODH component from *Ca*. E. thermophilum. Similarly, the experimentally determined molecular weight of 174-253 kDa for the Fmd/Fwd complex also appears to be incoherent with the previously described complexes (18, 19, 21). The compositions of both atypical complexes were elucidated through crystallographic snapshots.

The structure of the CODH was refined to a 1.89-Å resolution (Fig. 2A, Table S2). As reflected by the denaturing PAGE profile (Fig. 1B), the complex organizes as a dimer of three subunits: the α and ε subunits (respectively CAD7772032 and CAD7772037) already described in methanogens and an additional subunit homologous to the F_420_H_2_-oxidase domain of the sulfite reductase from *Methanothermococcus thermolithotrophicus* (Fig. 2A, Fig. S2, S3 and S4). Because of its tight binding on the CODH and its presence in the ACDS operon, this additional subunit will be referred to as ζ subunit (CAD7772047).

**Figure 2.**
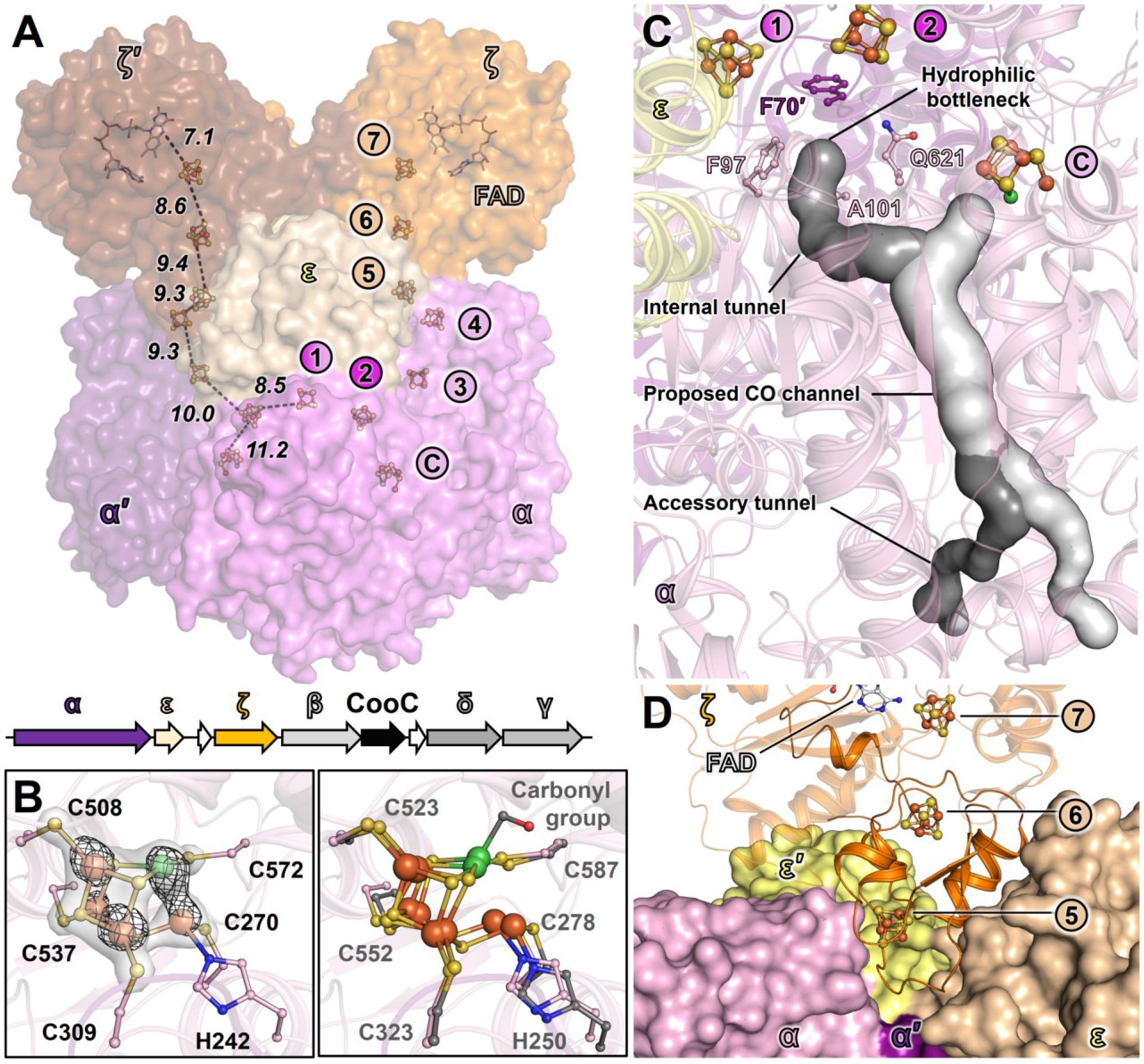
ACDS subcomplex from *Ca*. E. thermophilum. **A**. Overall structure of the subcomplex and its organization in the genome. The structure is shown as a surface. The distances between electron-transferring (metallo)cofactors in the α′ε′ζ′ half-complex are presented by dashed black lines and given in Å. The genomic organization of the genes encoding the complex is shown, with arrows colored by subunits (white for unannotated, unrelated, or pseudogene) and size depending on the gene length. **B**. Architecture of the C-cluster (left) and superposition with the homolog from *M. barkeri* (PDB 3CF4, colored grey, right panel). The 2*F*_o_-*F*_c_ and anomalous maps (collected at 12.67 keV), contoured at 3 and 5 σ, are shown as transparent white surface and black mesh, respectively. The carbonyl group modeled on the C-cluster of *M. barkeri* is absent in the structure of the ethanotroph. **C**. The different tunneling systems in α_2_ε_2_ζ_2_ structure predicted by the CAVER program are shown as surfaces and colored by tunnels (structuring residues in Fig. S7). **D**. ζ_2_ dimer (cartoon) bound on the α_2_ε_2_ core (surface). The ferredoxin-like N-terminal domain of the ζ subunit (1-83) is shown as a non-transparent cartoon. The ζ′ subunit has been omitted for clarity. **A-D**. Cofactors and residues are represented as balls and sticks with oxygen, nitrogen, sulfur, phosphorus, iron, and nickel colored in red, blue, yellow, light orange, orange and green, respectively. Carbons are colored according to the respective chains and white for the FAD.

The α_2_ε_2_ core of the complex can be reliably superposed on the homologous structure from *Methanosarcina barkeri* (Fig. S3A and B). The complete chains could be modeled except few residues at the C-terminal extremities of the α/α’ subunits and the N-terminal extension of the α subunit (1-30) conserved in archaea, which probably stabilizes the interaction with the β subunit (Fig. S5). While the α subunit exhibits a high structural conservation, the ε subunit slightly differs between *Ca*. E. thermophilum and *M. barkeri* due to local reorganizations to accommodate the ζ subunit (Fig. S3C). All metallo-cofactors harbored on the α subunit are coordinated by residues similar to those described in the archaeal homolog (Fig. S6). However, some substitutions in their close environments might tune their redox potentials (Fig. S6). The C-cluster operating the CO-oxidation is in a not-carbonylated reduced state with the CO site vacant in both α subunits occupying the asymmetric unit (Fig. 2B). In contrast, the C-cluster in *M. barkeri* has been found to harbor a CO-bound ligand, explaining the differences in geometry in the overlay (Fig. 2B). Because of their gaseous and hydrophobic natures, the substrate CO and product CO_2_ must transit through an internal hydrophobic channeling network that was experimentally identified in the structure of *M. barkeri* (16). Computational analyses of the internal cavities confirmed the conservation of this channeling system in the α subunit of the complex from *Ca*. E. thermophilum (Fig. 2C and S7). A main tunnel emanating from the C-cluster and ending up in two tunnels reaching the surface of the protein was detected and is proposed to be the CO-channel, as observed in the protein from *M. barkeri* and the bacterial homolog *Clostridium autoethanogenum* (25) (Fig. 2C and S7). Another internal cavity exhibiting ramifications within the α subunits has also been detected, but the diffusion of the CO_2_ and CO from the C-cluster in this extended channeling system is improbable due to a hydrophilic bottleneck restricting its access (Fig. 2C).

The ζ subunit, positioned at the intersection of α and ε subunits (Fig. 2D), is formed by a ferredoxin domain in its N-terminal part (1-83), an F_420_-reductase domain (84-349), and a C-terminal extension promoting the homodimeric interface (350-370). The F_420_-reductase domain shares high structural similarity with the F_420_-reducing subunit (FrhB) of the F_420_-reducing hydrogenase from *Methanothermobacter marburgensis* (*Mm*Frh, PDB 4OMF (26), Fig. S4A) and the F_420_H_2_-oxidizing sulfite reductase Fsr (Fig. S4A, (27)) with a comparable position and coordination of their (metallo)-cofactors (Fig. S8 and S9). The ζ subunits are anchored to the α_2_ε_2_ core through the ferredoxin domains, providing an attractive template to understand how soluble ferredoxins would dock on the archaeal ACDS complex (Fig. 2D). The ferredoxin domain acts as an electron bridge to electronically connect the C-cluster to the FAD site (Fig. 2A), which would allow the coupling of the CO-oxidation to F_420_-reduction.

The Fmd/Fwd complex was predicted to catalyze the second CO_2_-generating step occurring in ethanotrophy (7, 8). These CO_2_-releasing complexes have not been structurally unveiled yet, and their reaction mechanisms have been proposed to be analogous to the CO_2_-fixing systems from hydrogenotrophic methanogens for which structures are available (18, 19, 21, 28). In this scenario, the binuclear [Zn-Zn] center of the A subunit would hydrolyze the formyl-MFR into MFR and formate (or formic acid), this latter diffusing in an internal polar tunnel to the tungsten/molybdopterin site of the BD subunits to be oxidized into CO_2_ with the concomitant reduction of ferredoxins (Fig. 1A).

The structure of the Fmd/Fwd complex from *Ca*. E. thermophilum was refined at a resolution of 1.97-Å (Fig. 3A, Table S2, Fig S10). Compared to the initial prediction of a molybdenum-containing enzyme (8), the anomalous data confirmed the presence of a tungsten atom in the active site, therefore renaming the complex Fwd (Fig. 3B). The Fwd complex forms a heterohexameric assembly in which the canonical polyferredoxin FwdF subunit is replaced by an F_420_-reductase (CAD7775209, named FwdI, Fig. 3A). FwdI shares a similar organization and (metallo)-cofactor content to the ζ subunit of ACDS, with the exception of the absence of the C-terminal extension involved in dimerization (Fig. S4, S8-S10). The lack of the polyferredoxin FwdF is coherent with the apparent absence of higher organization in the crude extract (Fig. 1C), as FwdF acts as an electron bridge generating the multimerization of the Fmd/Fwd complexes and the establishment of larger enzymatic complexes (18, 19, 21). The six [4Fe-4S] clusters dispatched in FwdBGI would allow an efficient electron transfer between the tungstopterin center to the FAD, therefore possibly coupling formate oxidation to F_420_-reduction.

**Figure 3.**
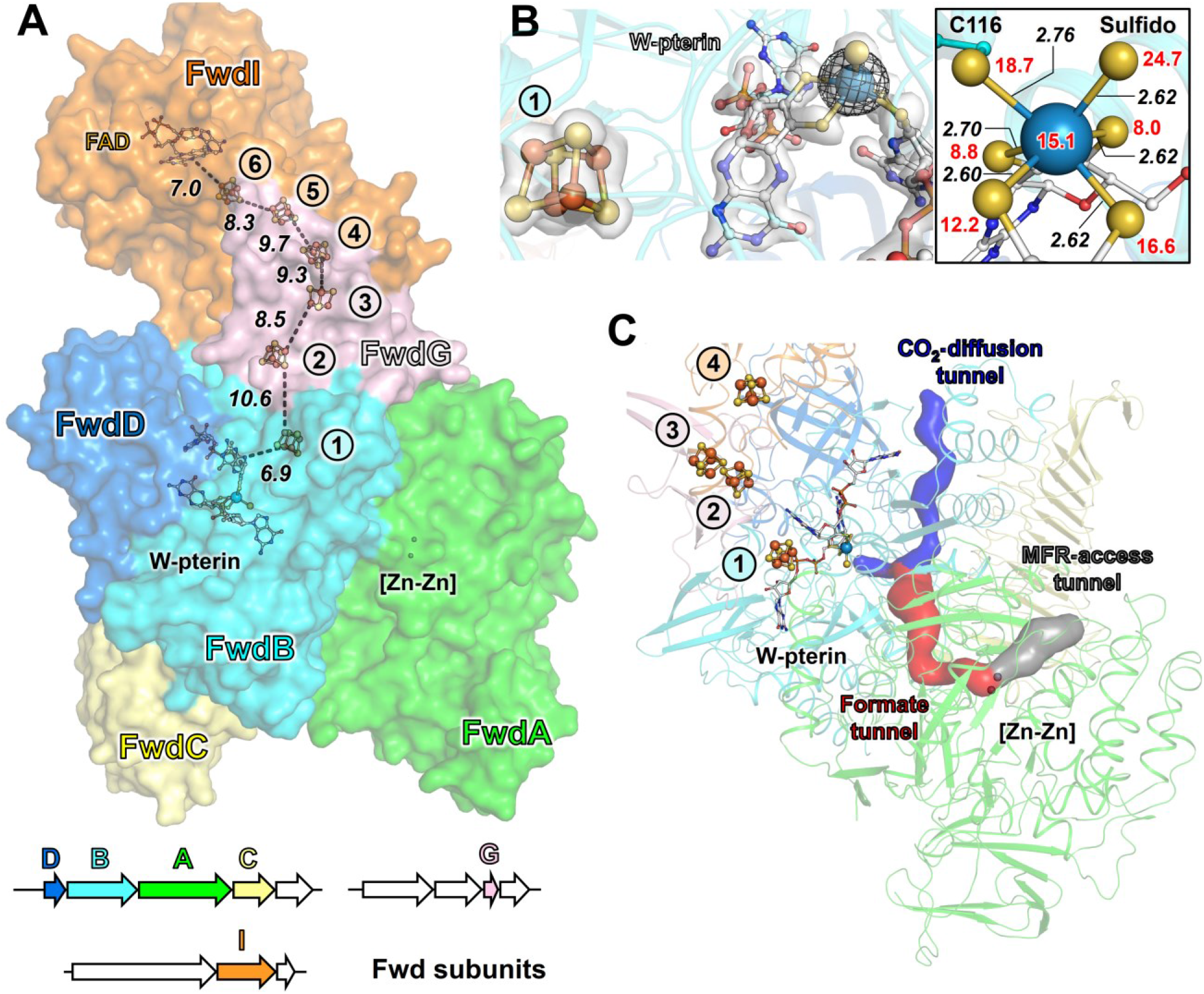
Fwd complex from *Ca*. E. thermophilum. **A**. Overall structure of the Fwd complex and its organization in the genome. The proteins are represented as surfaces with the distance between electron-transferring cofactors highlighted by dashed black lines and given in Å. The genomic organization of the genes encoding the complex is shown, with arrows colored by subunits (white for unannotated, unrelated, or pseudogene) and size depending on the gene length. **B**. Left, details of the tungstopterin site. The 2*F*_o_-*F*_c_ (1.5 σ) and anomalous map (7.0 σ, collected at ∼12.34 keV) are shown as white transparent surface and black mesh, respectively. Right, close-up of the tungsten ligands. Distances between the W atom and ligands are given in Å (black italic), and b-factors of atoms are given (red). **C**. Tunneling system in the Fwd complex from *Ca*. E. thermophilum. The tunnels predicted by the CAVER program are represented as surfaces and colored based on their proposed function. In all panels, the A, B, C, D, G, and I subunits are colored green, cyan, light yellow, marine blue, light pink, and orange, respectively. Cofactors and residues are represented as balls and sticks with oxygen, nitrogen, sulfur, phosphorus, iron, zinc, and tungsten colored in red, blue, yellow, light orange, orange, light grey, and blue-grey, respectively. Cofactors carbons are colored white.

The FwdABCDG core is similar to the structures of the formylmethanofuran dehydrogenase complex and subcomplex from *Methanothermobacter wolfei* (18) and *Methanospirillum hungatei* (21) (Fig. S10B). Both active sites, containing the metallocenters [Zn-Zn] and tungstopterin, are also highly structurally conserved (Fig. S11 and S12). The internal tunneling systems required for the transit of formyl-MFR, formate, and CO_2_ can be predicted from the structure (Fig. 3C, Fig. S13). Such conservation argues that the directionality of the overall reaction is not due to modifications of the metallo-cofactor, its coordination, or the active site architecture but is rather under the control of metabolic fluxes, including the final electron acceptor of the reaction.

The structural data gathered on the ACDS and Fwd complexes isolated from *Ca*. E. thermophilum points out that both systems would perform F_420_-reduction due to the acquisition of a functional module. Accordingly, we demonstrated that both enzymes use F_420_ as an electron acceptor, validating the functional assembly observed in the crystal structures (Fig. 4A and B). These overall reactions would be largely exergonic when considering the standard midpoint redox potentials of the CO-oxidation (*E*_0_′ CO/CO_2_ = -520 mV (29)), formyl-MFR oxidation (*E*_0_′ formyl-MFR/MFR + CO_2_ = -530 mV (30)) coupled to F_420_-reduction (*E*_0_′ F_420_H_2_/F_420_ = -340 mV (31)). The highly favorable coupling could represent the “thermodynamic pull” of anaerobic ethane oxidation, preventing the reversal of the pathway in the absence of ethane (i.e., CO_2_ fixation dependent on F_420_H_2_-oxidation would be improbable under physiological conditions). This might be particularly important under common seafloor conditions, where concentrations of CO_2_ largely exceed those of ethane. On the same line of thoughts, the exergonic process could counterbalance one or several unfavorable enzymatic reactions occurring during the uncharacterized conversion of ethyl-CoM into acetyl-CoA, explaining why the ζ and FwdI subunits are apparently conserved in the other cultured ethanotroph *Ca*. A. ethanivorans ((7), Fig. S14 and S15). In comparison, it has been assumed that methanogens, methanotrophs, and longer-chain alkanotrophs (using the β-oxidation pathway (11-13)) depend on ferredoxin for the CO_2_-releasing steps. A comparison of the ACDS and Fwd/Fmd operons organization indicates that methanogens and alkanotrophs do not seem to have the genetic capability of coupling the CO_2_-releasing steps to F_420_-reduction except for a few exceptions detailed in supplementary (Fig. S14 and S15). Accordingly, the activities of F_420_-reduction coupled to CO or furfurylformamide oxidation could not be detected in the cell extracts of ANME-1 and ANME-2 or the *Methanosarcinales M. barkeri* grown on either acetate or methanol (Fig. 4C). It can be hypothesized that the second CODH isoform which could include a ζ subunit found in other microorganisms such as some ANME-1 species might be involved in CO-detoxification and further studies will have to clarify the roles of these proteins.

**Figure 4.**
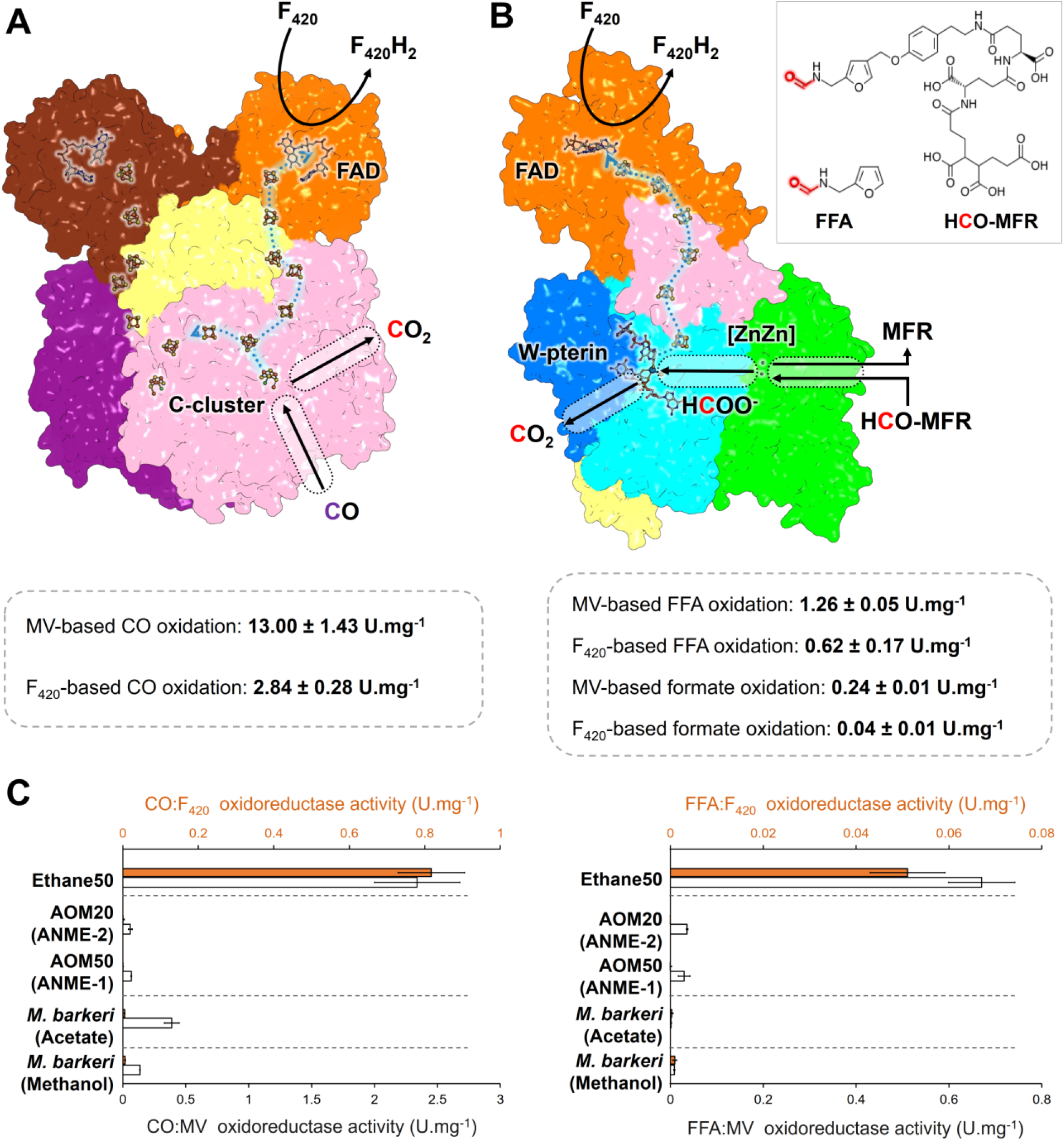
Overall reactions and activity comparison of both complexes in cell extracts from ethanotroph, methanotrophs, and methanogens. **A and B**. Proposed reactions and activity measurements for CODH (A) and Fwd (B) complexes. Acetate (100 mM) was not used as a substrate by the Fwd complex with either MV or F_420_. The chemical structure of formylmethanofuran (MFR) is the one described in *Methanothermobacter thermautotrophicus* (34). **C**. Activity measurements performed in soluble extracts from the ethanogen, methanotrophs, and *M. barkeri* during aceticlastic and methylotrophic methanogenesis. **A-C**. Activities are given in μmol of substrates oxidized (CO, FFA or formate) per minute per mg of pure enzyme or soluble proteins.

The genome of ethanotrophs lacks ferredoxin-dependent systems such as Rnf or Ech, suggesting that F_420_ is a central electron carrier in ethanotrophy. In our proposed model (Fig. 5), the highly expressed Fpo complex (Table S4) would be the only energy-conserving system that would allow ion translocation across the membrane to fuel the ATP synthase. The electrons extracted from the F_420_H_2_ pool by Fpo would be consumed through the thermodynamically favorable sulfate reduction pathway of the bacterial partner. The transfer of electrons from the archaeal Fpo to the bacterial quinones would be operated through an elusive path that might imply conductive nanowires (8). The cytoplasmic pool of F_420_H_2_ will be replenished by the ethanotrophic catabolism. The stoichiometry of the ethane/sulfate oxidoreduction performed by the consortium (4 moles of ethane oxidized for 7 moles of sulfate reduced (8)) indicates that a total of seven F_420_H_2_ could be potentially obtained from the complete oxidation of one ethane molecule. The oxidation of acetyl-CoA by the ACDS and the reactions occurring in reverse methanogenesis (by the Fwd complex and the methylenetetrahydromethanopterin dehydrogenase and reductase) would reduce four out of seven F_420_. We propose that the missing reduced F_420_s are derived from the two oxidative steps occurring in the metabolic transformation of ethyl-CoM to acetyl-CoA. In our hypothesis, one F_420_H_2_ and one reduced ferredoxin would be produced, and the ferredoxin will be oxidized concomitantly with the heterodisulfide CoM-S-S-CoB by the highly expressed F_420_H_2_-oxidizing electron-bifurcating Hdr to generate two F_420_H_2_ (Fig. 5 and Table S4).

**Figure 5.**
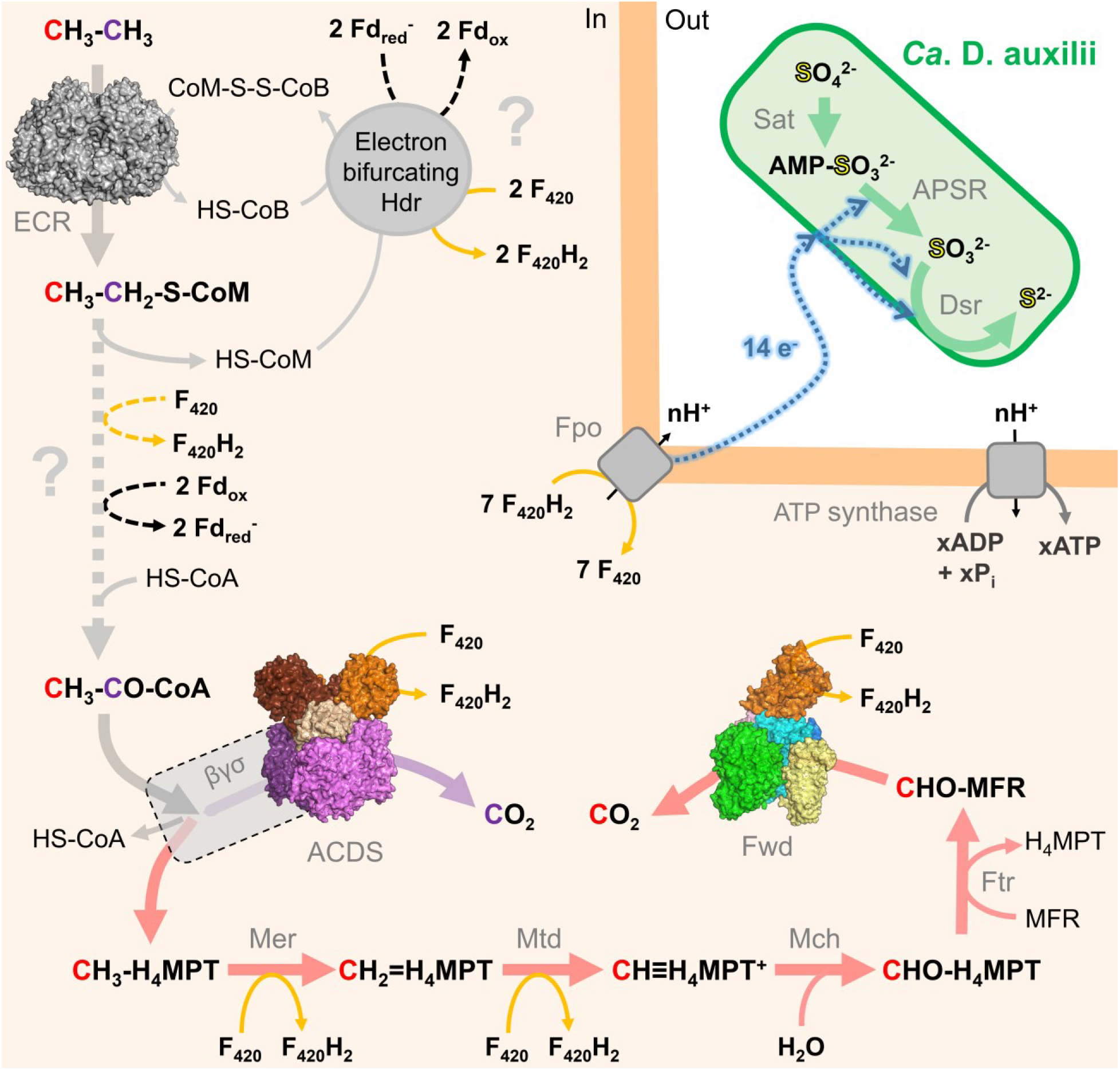
Proposed catabolic metabolism in the Ethane50 consortium. The structurally characterized enzymes are shown as surface representations. The catabolic reactions are presented as large arrows colored in grey, purple, red, and green, corresponding to the C2 part of ethanotrophy, ACDS activity, reverse methanogenesis, and sulfate reduction, respectively. A large dashed arrow indicates the yet uncharacterized ethyl-CoM to acetyl-CoA conversion. Orange arrows indicate F_420_ reduction or F_420_H_2_ oxidation events. A question mark highlights the uncharacterized reactions that would employ ferredoxin. The interspecies electron transfer is schematized in a blue dashed line, and the transfer mechanism was omitted in the figure for clarity. The exact number of ions translocated by the Fpo system is not known and is therefore labelled “n” and hypothesized to be protons. The ion/ATP ratio of the ATP synthase is also not known, and therefore “x” ATP is produced, while ions are proposed to be protons. The stoichiometry of sulfate reduction is not respected.

The physiological utilization of the F_420_H_2_ pool can be extended to assimilatory (i.e., in nitrogen assimilation through the putative F_420_-dependent glutamate synthase) and anabolic pathway, as suggested by the numerous *frhB* homologs in the genomes of ethanotrophs (Fig. S15). This unexplored reservoir of reactions coupled to F_420_(H_2_) oxidoreduction must contain potential unknown metabolic routes and, among them, the reactions behind the ethyl-CoM transformation that remains to be elucidated.

By isolating native systems from a thermophilic microbial enrichment, this study revealed new pieces of the molecular puzzle of anaerobic ethane oxidation. The structural knowledge acquired provides a reasonable model of the docking sites of the ferredoxin in the ACDS and Fwd/Fmd complexes in other anaerobes and presents another remarkable example of surface remodeling to allow an electrical connection. The catalytic sites and channeling systems in the two complexes also argue that the reversibility of the CO_2_-fixation/generation reactions is not tuned by molecular determinants such as cofactor modifications and substitutions but rather dictated by the cellular metabolic fluxes. Importantly, we discovered that the ethanotroph *Ca*. E. thermophilum relies on F_420_ instead of ferredoxin as an electron acceptor for CO_2_-generating enzymes by acquiring F_420_-reducing modules. By taking such a different metabolic route compared to what has been learned in methanogens, the ethanotrophs would benefit from these exergonic reactions to drive ethane oxidation, with bacterial sulfate reduction as the final electron sink. Applying the native approach described in this work to ANMEs or other alkanotrophs enrichments will progressively unveil the global picture of how microbial Life can derive cellular energy from alkane transformation and which specific strategies have been applied by these astonishing microbes to optimize their catabolisms.

## Acknowledgments

We are deeply thankful to Cedric J. Hahn for his help in the culture of the enrichment and during enzyme purification. We thank the Max Planck Institute for Marine Microbiology and the Max Planck Society for their continuous support. We thank the SOLEIL and Swiss Light Source (SLS) synchrotrons for beam time allocation and the respective beamline staffs of PROXIMA-1 and X06DA for assistance with data collection. We also acknowledge Christina Probian, Ramona Appel, and Mélissa Belhamri for their invaluable support in the Microbial Metabolism research group.

## Author contributions

O.N.L., G.W., and T.W. designed the research. O.N.L., and G.W. performed cultivation and culture experiments. O.N.L., and T.W. purified and crystallized the proteins. O.N.L., and T.W. collected X-ray data and built the models. O.N.L. and T.W. analyzed the structures. O.N.L. performed the activity measurements. O.N.L., and T.W. interpreted the data and wrote the paper, with contributions and final approval of all co-authors.

## Data and materials availability

All structures were validated and deposited in the Protein DataBank (PDB) under the following accession numbers: 8RIU, Crystal structure of the F_420_-reducing carbon monoxide dehydrogenase component and 8RJA, Crystal structure of the F_420_-reducing formylmethanofuran dehydrogenase complex. All other data are available in the manuscript or the supplementary materials.

