## Supplementary information for "F_420_ reduction as a cellular driver for anaerobic ethanotrophy"

**This PDF file includes:**

Materials and Methods
Figs. S1 to S15
Tables S1 to S4

### Materials and Methods

**Origin and cultivation of Ethane50, AOM50, and AOM20 enrichments.** The Ethane50 enrichment was used for enzyme purification derived from heated sediments of the Guaymas Basin hydrothermal vents sampled during RV Atlantis mission AT 37-06 with submarine Alvin in December 2016. The culture conditions were previously described (1), and the studied biomass is similar to that used for the purification of the ethyl-coenzyme M reductase (ECR), previously characterized (2). The AOM50 culture was derived from the Guaymas Basin (sampled during RV Atlantis mission AT 15-56 in December 2009). The culture conditions and microbial composition were described before (3-6). The AOM20 culture, dominated by ANME-2, is derived from sediments from Amon Mud Volcano, Eastern Mediterranean Sea, retrieved during Nautinil Expedition RV Atalante in 2003. Cultivation was performed at room temperature (20 °C) with methane as sole energy substrate. The microbial composition of this culture is described in Wegener *et al.* (2016) (7).

**Cultivation of methanogenic archaea.** *Methanosarcina barkeri* DSM 800 was obtained from the Deutsche Sammlung von Mikroorganismen und Zellkulturen GmbH, Braunschweig, Germany and was cultivated at 37 °C under strict anaerobic conditions in a medium whose composition was already described (8). Sterile and anoxic methanol (1 % v/v) or 100 mM sodium acetate was added as carbon and energy source. The initial gas phase contained $N_2/CO_2$ (90:10 %) at 50 kPa. The overpressure coming from the metabolic activity during growth was regularly evacuated. The cells were harvested in the late-exponential phase by centrifugation for 30 minutes at 17,000*g* and kept frozen at -80 °C under anaerobic conditions before use.

**Protein extraction and purification.** Cells collected from exponentially growing Ethane50 culture were used for protein extraction. The medium was removed with a stainless steel needle by applying an overpressure of $N_2/CO_2$ (with a 90:10 % ratio). After a 3 min-long flushing with $N_2/CO_2$ (90:10 %), the cells were pelleted by centrifugation for 15 min at 16,250*g* in an anaerobic chamber filled with an $N_2/CO_2$ atmosphere (90:10 %) at room temperature, and the supernatant was removed. Cells were suspended in SR medium (9) and stored at −80 °C under an $N_2/CO_2$ (90:10 %) atmosphere until purification. Because of black aggregate precipitates, the biological quantities of the sample could not be determined.

Because of the extremely limited biomass, only two purification procedures could have been performed and were used for this work. Only one is described in detail. Cell lysis and preparation of extracts were performed in an anaerobic chamber filled with an $N_2/CO_2$ atmosphere (90:10 %) at room temperature. A volume of 15 ml of sedimented cells was suspended in 16 ml of 50 mM tricine/NaOH buffer pH 8.0, 2 mM dithiothreitol (DTT; buffer A). The lysis protocol included a sonication step (BANDELIN Sonopuls HD 2200) followed by five rounds of French Press at around 1,000 PSI (6.895 MPa), yielding a homogenous deep-black extract. The French press cell was flushed with $N_2$ and washed twice with anoxic buffer A before use. Soluble extract was obtained by ultracentrifugation at 140,000*g* for one hour at 4 °C. Enzymes purification was carried out under anaerobic conditions in a Coy tent, filled with an $N_2/H_2$ atmosphere (97:3 %), at 20 °C and under yellow light. For each step, chromatography columns were washed with at least three-column volumes (CV) with the corresponding loading buffer, and samples were filtrated on 0.2 µm filters prior to loading. During purification, the enzymes were followed by high-resolution clear native polyacrylamide gel electrophoresis (hrCN PAGE, see below), sodium dodecyl sulfate PAGE (SDS PAGE), and absorbance monitoring at 280, 415 and 550 nm.

Extracts were diluted with buffer A to obtain a final 15-fold dilution before being loaded on 4 × 5 ml anion exchanger HiTrap™ Q HP columns (GE Healthcare) equilibrated with the same buffer. After a 2 CV washing, proteins were eluted with a 0 to 0.40 M NaCl linear gradient for 6 CV at a 1 $ml.min^{-1}$ flow rate. The fractions containing both the CO-dehydrogenase (CODH) and the formylmethanofuran dehydrogenase (Fwd) complexes from *Ca*. E. thermophilum eluted between 0.38 M and 0.40 M NaCl. The pooled fractions were diluted with 2 volumes of an anoxic 50 mM Tris/HCl buffer pH 7.6, 2 mM DTT (buffer B) containing 2 M ammonium sulfate, before being loaded on a Source™ 15PHE 4.6/100 PE (GE Healthcare) equilibrated with the same buffer. After washing, proteins were eluted with a 1.60 to 0 M ammonium sulfate linear gradient for 53 CV at a 1 $ml.min^{-1}$ flow rate. The CODH complex eluted between 1.13 M and 0.80 M ammonium sulfate. In the other purification, the hydrophobic exchange chromatography step using the Source™ 15PHE 4.6/100 PE column has been performed twice to enhance sample purity (see Fig. 1B and Table S1). Fractions of interest were pooled, concentrated on a 30-kDa cut-off centrifugal concentrator (nitrocellulose, Vivaspin from Sartorius) and injected on a Superdex 200 Increase 10/300 GL. The size-exclusion chromatography was performed in 25 mM Tris/HCl buffer pH 7.6,

10 % (v/v) glycerol, 2 mM DTT (buffer C) at a 0.4 $ml.min^{-1}$ flow rate. The CODH complex eluted in a Gaussian peak with an 11.04 ml elution volume. The protein was directly used for anaerobic crystallization and activity measurements. The Fwd complex eluted between 1.50 M and 1.13 M ammonium sulfate during hydrophobic exchange chromatography. Fractions of interest were pooled and concentrated on a 30-kDa cut-off centrifugal concentrator (nitrocellulose, Vivaspin from Sartorius), and the buffer was exchanged for buffer C. The protein was directly used for anaerobic crystallization and enzymatic activities.

Protein concentration was estimated by the Bradford method (Bio-Rad Laboratories, Munich, Germany) for all samples. A bovine serum albumin (BSA) standard was used to estimate protein concentration.

**Preparation of cell extracts.** The different extracts used for activity measurements presented on Figure 4 were prepared in an anaerobic chamber filled with an $N_2/CO_2$ atmosphere (90:10 %) at room temperature. The extracts from *M. barkeri* were obtained from anaerobically frozen pellets of 4.24 g and 1.15 g (wet weight) of cells grown on methanol and acetate, respectively. The biomass used for the preparation of the extracts from ANMEs was obtained from the sedimentation of the cultures of AOM20 and AOM50 (around 1.5 ml and 2 ml sedimented cells, respectively). For all organisms, the biomass was diluted with 5 ml of buffer B before a sonication step (BANDELIN Sonopuls HD 2200). Five rounds of French Press at around 1,000 PSI (6.895 MPa) were used as an additional lysis step. Unbroken cells and debris were removed by ultracentrifugation at 140,000*g* for one hour at 4 °C. Protein concentration was estimated by the Bradford method using BSA as standard. The protein concentration from the ANMEs extracts was increased by using a 10-kDa cut-off centrifugal concentrator (nitrocellulose, Vivaspin from Sartorius).

**High-resolution clear native polyacrylamide gel electrophoresis (hrCN PAGE).** The hrCN PAGE protocol was adapted from Lemaire *et al.* (2018) (10). The electrophoresis was performed in an anaerobic chamber filled with a $N_2/CO_2$ (90:10 %) atmosphere. Fresh anaerobic samples were used. Glycerol (20 % v/v final) was added to samples, and 0.001 % (w/v) Ponceau S was used as a protein migration marker. The anaerobic electrophoresis cathode buffer contained a buffer mixture of 50 mM tricine/NaOH, 15 mM Bis-Tris at a pH 7.0 supplemented with 0.05 % (w/v) sodium deoxycholate, 0.01 % (w/v) dodecyl maltoside and 2 mM DTT. The anaerobic anode buffer contained 50 mM Bis-Tris buffer, pH 7.0, and 2 mM DTT. hrCN PAGEs were carried out using an 8 to 15 % linear polyacrylamide gradient, incubated overnight in an anaerobic tent, and soaked in an anaerobic cathode buffer. Gels were run with a constant 20 mA current using a PowerPac™ Basic Power Supply (Bio-Rad). After electrophoresis, gels were stained with Instant Blue™ (Expedeon) or used for enzymatic staining (see below). Estimation of complexes size on gels was performed by measuring the distance between the center of the sample wells and the center of the protein bands. The protein ladder (NativeMark unstained protein ladder, Fischer Scientific) was used to establish the standard curve. The equation derived from the standard curve was used to estimate the size of standards and CODH or Fwd complexes.

**Determination of the molecular weight and oligomeric state by size-exclusion chromatography.** The chromatography was performed on a Superdex 200 Increase 10/300 GL (GE Healthcare, Munich, Germany) in 25 mM Tris/HCl pH 7.6, 2 mM DTT, 10% (v/v) glycerol at a 0.4 ml.min$^{-1}$ flow rate and in an anaerobic Coy tent containing an $N_2$/$H_2$ (97:3%) atmosphere. The elution volume determined from a high molecular weight range gel filtration calibration kit (GE Healthcare, Munich, Germany) was used to establish the protein standard curve. Estimations of the molecular weight from standards and CODH or Fwd complexes were derived from the equation of the standard curve.

**Enzymatic assays.** The enzymatic activity measurements performed during purification and on purified enzymes were performed in 50 mM Tris/HCl pH 7.6, 2 mM DTT, under anaerobic conditions and at 50 °C. A final concentration of 5 mM methyl-viologene (MV) was used, its reduction being spectrophotometrically followed at 600 nm on a Carry 60 spectrophotometer in sealed quartz cuvettes. A molecular extinction coefficient of 14,181 M$^{-1}$.cm$^{-1}$ was experimentally determined under these conditions and used for calculations. The presented activities are in µmol of oxidized substrate (CO, furfurylformamide or formate) per min per mg of protein, considering that 2 moles of MV are reduced per mole of the oxidized substrate. For CODH activity, the reaction was initiated by the addition of 0.2 ml of 100 % CO in an anaerobic quartz cuvette of 1.4 ml containing 1 ml of reaction mixture. For furfurylformamide/formate dehydrogenase activity

measurements, activity was started by the addition of furfurylformamide (10 mM final) or formate (10 or 100 mM final) solutions prepared in anoxic deionized water. No Fwd activity could be detected with either 10 or 100 mM acetate. The $F_{420}$ reduction by both enzymes was monitored in similar conditions, using 18.8 µM of $F_{420}$ purified from *M. thermolithotrophicus*. F420 was prepared by following the previously published protocol (11). $F_{420}$ reduction was monitored at 420 nm, using an experimentally determined molecular extinction coefficient of 41,503 $M^{-1}.cm^{-1}$. The presented activities are in µmol of oxidized substrate per min per mg of protein, considering that a mole of $F_{420}$ is reduced per mole of the oxidized substrate. 50 µM of flavin adenine dinucleotide (FAD) and flavin mononucleotide (FMN) was added to the enzymes stock before activity measurement to warrant flavin saturation and maximal activity of the $F_{420}$-reducing subunits. Final concentration of FAD and FMN during activity measurement was inferior or equal to 0.1 µM. addition of FAD and FMN did not affect measurements in the absence of the enzymes. Activity in crude extracts was monitored in similar conditions, except that FAD and FMN concentration was 100 µM. Final concentration of FAD and FMN during activity measurement was inferior or equal to 1.2 µM. addition of FAD and FMN did not affect measurements in the absence of the extracts. Experiments were run at least in triplicates. Activities measured in the extracts of *M. barkeri*, AOM50, AOM20, and Ethane50 were performed at 37, 50, 20 and 50 °C, respectively. The indicated units refer to µmole of CO or furfurylformamide oxidized per minute per mg of protein, using the parameters and coefficients listed above.
In-gel viologen-based activity staining was performed as previously described (12). Shortly, activity staining was performed in 10 ml of anoxic 50 mM Tris/HCl buffer pH 7.6, 2 mM DTT, 5 mM MV and 1 mM of 2,3,5-triphenyltetrazolium chloride (TTZ). The latter component was used to have an oxygen-insensitive staining. Native gels were loaded with 10, 5, or 2 µg of extracts or pure enzymes. After electrophoresis, gels were transferred to a 0.5 l Duran bottle containing the staining solution described above. The gas phase was changed for 100 % $N_2$ at 50 kPa. For the CODH activity, 20 ml of 100 % CO was added. Formate (10 or 100 mM final) or furfurylformamide (10 mM final) was added for Fwd activity. The reaction was started by the addition of 1 ml of an anoxic solution of 50 mM MV and 10 mM TTZ. Reactions were performed at 50 °C and stopped by bottle opening under a fume hood and transfer in aerobic deionized water.

The equations of the monitored reactions are (FA standing for furfurylamine):

**MV-based CO oxidation:** $CO + H_2O + 2\ MV_{ox} \rightarrow CO_2 + 2\ H^+ + 2\ MV_{red}^-$

**$F_{420}$-based CO oxidation:** $CO + H_2O + F_{420} \rightarrow CO_2 + F_{420}H_2$

**MV-based FFA oxidation:** $FFA + H_2O + 2\ MV_{ox} \rightarrow CO_2 + 2\ H^+ + FA + 2\ MV_{red}^-$

**$F_{420}$-based FFA oxidation:** $FFA + H_2O + F_{420} \rightarrow CO_2 + FA + F_{420}H_2$

**MV-based formate oxidation:** $HCOO^- + 2\ MV_{ox} \rightarrow CO_2 + H^+ + 2\ MV_{red}^-$

**$F_{420}$-based formate oxidation:** $HCOO^- + H^+ + F_{420} \rightarrow CO_2 + F_{420}H_2$

**Protein crystallization.** All crystals were obtained at 20 °C in a Coy tent under an N2/H2 (97:4 %) by using the sitting drop method on a 96-Well MRC 2-Drop Crystallization Plates in polystyrene (SWISSCI). The crystallization reservoir contained 90 µl of mother liquor. Crystallization drops contained a mixture of 0.6 µl protein and 0.6 µl precipitant.

Crystals of the $\alpha_2\varepsilon_2\zeta_2$ ACDS subcomplex from *Ca.* E. thermophilum were obtained in 20 % (w/v) polyethylene glycol 6,000, 100 mM MES pH 6.0, and 200 mM ammonium chloride (JBS Wizard™ crystallization screen from Jena Bioscience). The initial protein concentration was 13.0 mg.ml$^{-1}$ and 1 mM final of both FAD and FMN was added to the protein before crystallization.

Crystals of the Fwd complex from *Ca.* E. thermophilum were obtained in 18 % (w/v) polyethylene glycol 8,000, 200 mM sodium acetate trihydrate and 100 mM sodium cacodylate, pH 6.5 (SG1™ crystallization screen from Molecular Dimensions). The initial protein concentration was 4.64 mg.ml$^{-1}$.

**X-ray data collection, model building, and refinement.** Crystals of the $\alpha_2\varepsilon_2\zeta_2$ ACDS subcomplex and the Fwd complex were soaked for a few seconds in the crystallization condition supplemented with 30 % (v/v) glycerol before being frozen in liquid nitrogen under anoxic conditions prior to X-ray diffraction studies. All diffraction experiments were performed at 100 K. The structure of the $\alpha_2\varepsilon_2\zeta_2$ ACDS subcomplex was initially solved by performing a single anomalous dispersion experiment at the Fe K-edge (Table S2) using the *SHELX* package (13), using diffraction data collected on the PXIII beamline (X06DA) from the Swiss Light Source (SLS, Villigen, Switzerland). Due to the relatively low resolution, these data were only used to experimentally solve the structure, and no extended model was built. The resolution was extended to 1.893-Å by a second dataset collected at a wavelength of 0.98 Å on the PROXIMA-1 beamline from SOLEIL (Paris-Saclay, France). The structure of the Fwd complex was solved by molecular replacement using the Fwd complex from *M. thermolithotrophicus* (PDB 5T5M (14)) as a template, using diffraction data collected on the PXIII beamline (X06DA) from the SLS. The data were processed and scaled with autoPROC (15). Crystallographic data presented anisotropy along the following axes: a=1.892 Å, b=2.155 Å, and c=2.301 Å for the $\alpha_2\varepsilon_2\zeta_2$-ACDS subcomplex and a=1.885 Å, b=2.291 Å, and c=3.018 Å for the Fwd complex. The data were accordingly further processed with *STARANISO* correction integrated with the *autoPROC* pipeline (16). All models were manually built via COOT (17) and refined with *PHENIX* (version 1.20.1-4487). The last refinement steps were performed by refining with a translation libration screw (TLS) and were validated by the MolProbity server (18) (http://molprobity.biochem.duke.edu). Both models were refined with hydrogens in the riding position. Hydrogens were omitted in the final deposited models. The PDB ID codes of the structures are 8RIU and 8RJA for the $\alpha_2\varepsilon_2\zeta_2$-ACDS subcomplex and the Fwd complex, respectively. Data collection and refinement statistics for the deposited models are listed in Table S2.

**Structural analyses.** All figures were generated and rendered with PyMOL (Version 2.2.0, Schrödinger, LLC, New York, NY, USA). Internal tunnels predictions were performed by the *CAVER* tool (19). The analysis of the $\alpha_2\varepsilon_2\zeta_2$ ACDS subcomplex was performed by applying a probe radius of 1.0 Å and starting from the Ni atom of the C-cluster. The analysis of the Fwd tunneling system was performed by applying a probe radius of 1.0 Å and starting from the sulfido ligand of the tungstopterin, ignoring the Zn atoms of the [Zn-Zn] binuclear center.

**Bioinformatic analyses.** The proteins used for model construction and phylogenetic analysis were extracted from the genome assemblies from *Ca*. E. thermophilum ( GenBank: LR991654.1, (1)), *Ca*. A. ethanivorans (GenBank: RPGO01000021.1, (20)), *Methanosarcina barkeri* MS (Genbank CP009528.1), *Methanothermobacter wolfei* (GenBank JAAYMZ010000098.1, (21)), ANME-1 isolates (GenBank QENH01000201.1 and PQXB01000001.1), *Candidatus* Methanoperedens nitroreducens (GenBank JAIOIS010000036.1, (22)), *Candidatus* Synthrophoarcheum butanivorans (GenBank LYOR01000001.1, (23)), *Candidatus* Synthrophoarcheum caldarius

(GenBank. LYOS01000001.1, (23)), *Candidatus* Methanoliparum thermophilum (GenBank RXIF01000004.1, (24)) and *Candidatus* Methanoliparum whitmanii (obtained from the NODE database under the ID OED248975 (25), available at https://www.biosino.org/node/analysis/detail/OEZ007026). The protein sequences and putative operon organizations were obtained from the genomes using the Operon Mapper webserver (26). Gene length was taken into consideration for the construction of figures. The proteins annotation of ACDS and Fwd/Fmd subunits are derived from the available structures or by a BLAST search against the predicted protein sequences using as query the sequences from the different subunits of the characterized ACDS from *M. barkeri* (27), Fwd complex from *M. wolfei* (14) and $F_{420}$-reducing subunit of the $F_{420}$-reducing hydrogenase (28). The presented percentages of identity and query coverages are extracted from the BLAST analysis. The other proteins were manually annotated by BLAST research in the PDB and SwissProt databanks. The phylogenetic tree was constructed with sequences of all proteins homologous to ACDS ζ subunit and FwdI using the maximum likelihood method and was generated with the MEGA program (29) by using an alignment constructed with *MUSCLE* (30). A total of 200 replicates were used to calculate node scores. Functions are derived from the available literature or by similarity with existing enzymes.

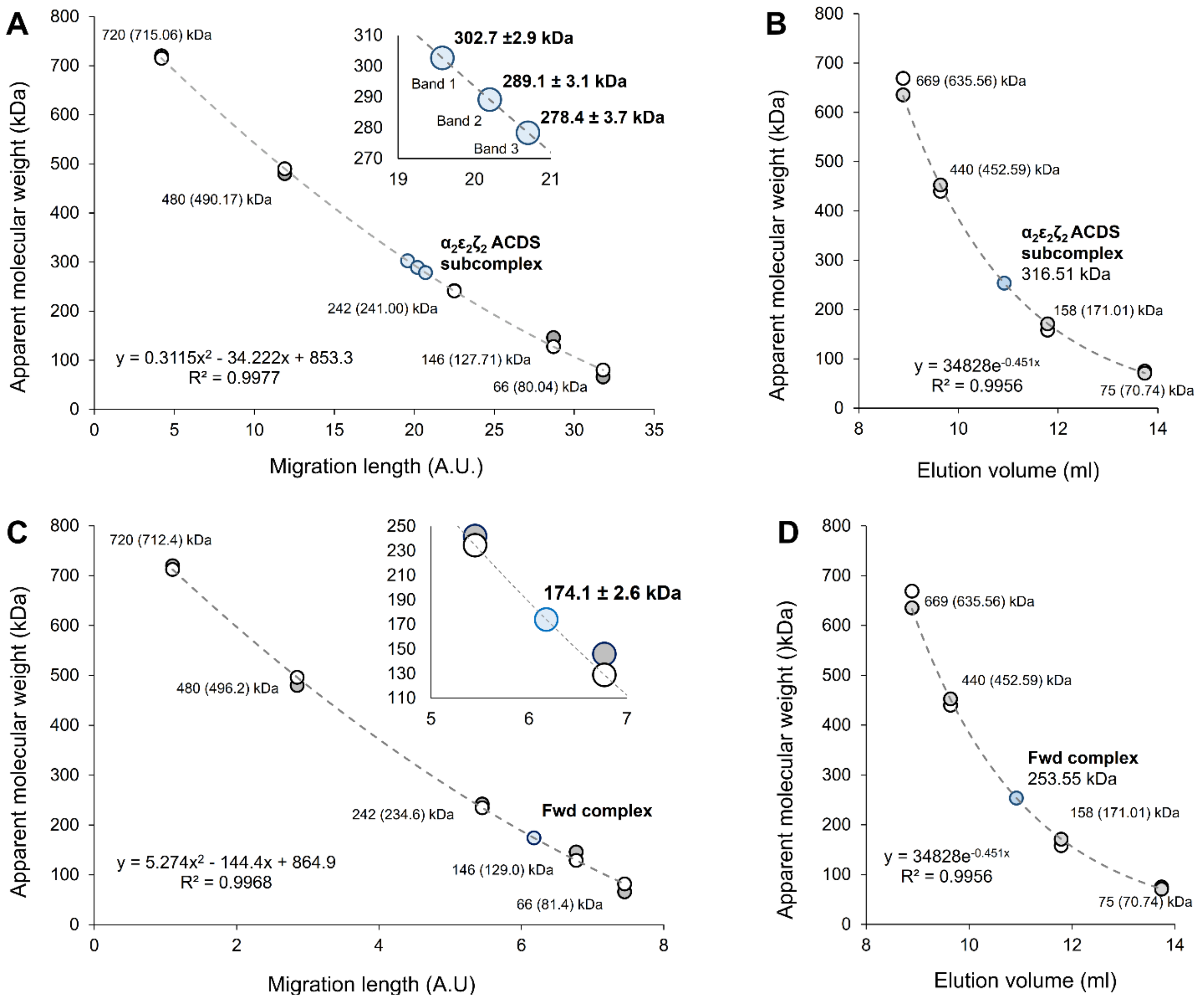

**Figure S1. Size estimation of the complexes purified from *Ca.* E. thermophilum on native PAGE and size exclusion chromatography. A and C.** Size estimation of the ACDS subcomplex (A) and Fwd complex (C) on native gels. The size and migration length of the proteins from the commercial ladder (prestained PageRuler from ThermoScientific, Germany) were used to calculate a fit (dashed grey line, equation, and $R^2$ indicated) to determine the molecular weights. White and grey dots correspond to the theoretical and measured molecular weights from the protein ladder. Blue dots display the molecular weight calculated from the fit for ACDS and Fwd enzymes. While the ACDS subcomplex appears to behave as a single population in the crude extract, the isolated enzyme seems to decompose in several populations, probably due to purification-dependent artifacts (e.g., partial loss of cofactors or stabilizing ions). The inset shows a close-up of the graph to describe the size of the different bands stained by CODH activity. **B and D.** Size determination of the ACDS subcomplex (B) and Fwd complex (D) by size-exclusion chromatography using elution volumes of standard proteins (High Molecular Weight range Gel Filtration Calibration Kit, GE Healthcare, Munich, Germany) and the purified enzymes. The size and elution volumes of the protein from the commercial ladder (white dots) were used to calculate a fit (dashed grey line, equation, and $R^2$ indicated), used for the determination of the theoretical size of the standard proteins (grey dots) and the proteins from *Ca.* E. thermophilum (blue dots).

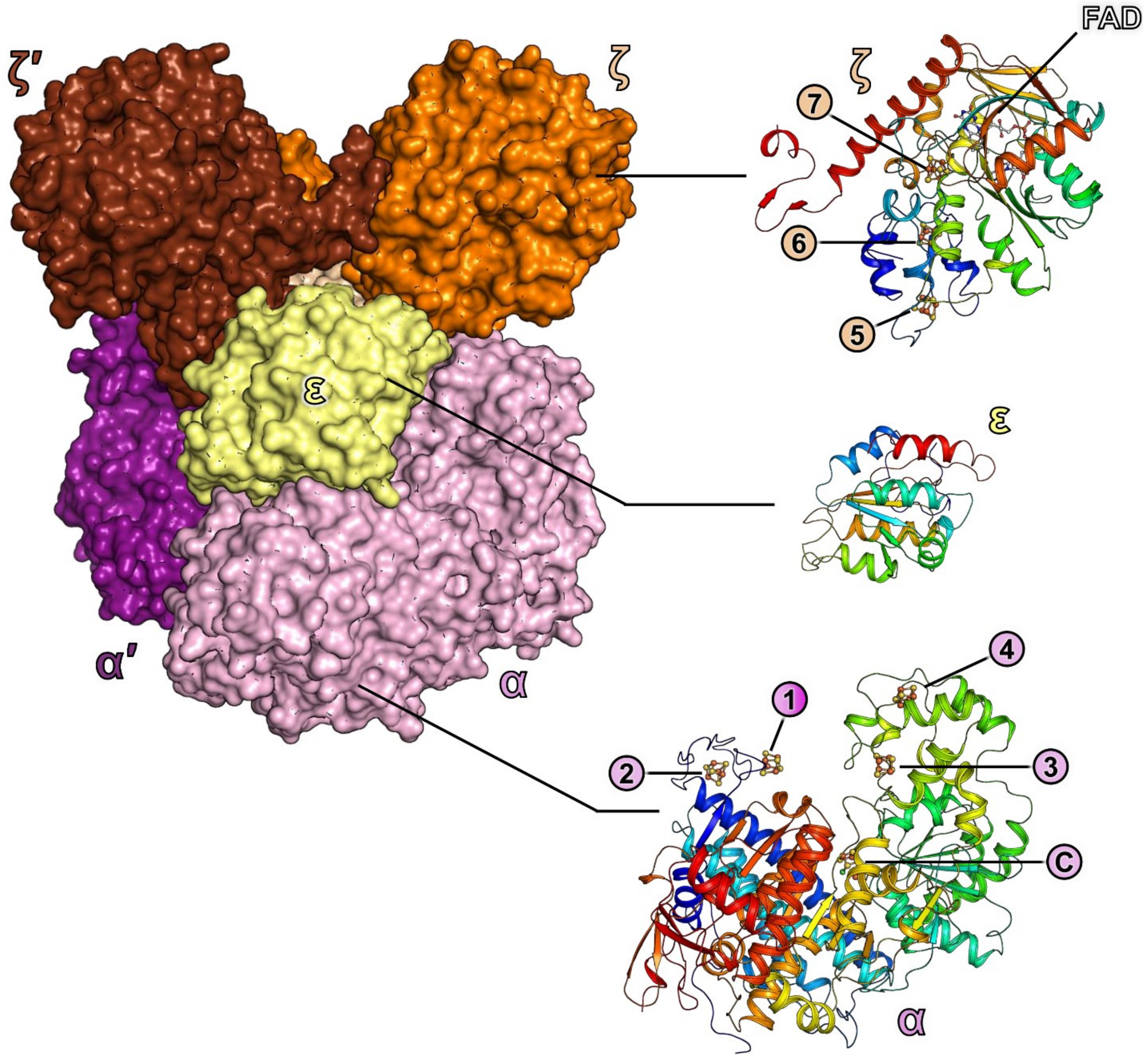

**Figure S2. Structure of the individual components of the $\alpha_2\varepsilon_2\zeta_2$ subcomplex from *Ca.* E. thermophilum.** The complex is shown in the upper left panel as a surface with α, ε, ζ, α′, ε′, and ζ′ subunits colored light pink, light yellow, orange, deep purple, wheat, and brown, respectively. Individual α, ε, and ζ subunits are presented in cartoons and colored in a rainbow from blue to red (N-terminus to C-terminus), in the same pose as the top left panel. Cofactors are displayed as balls and sticks and atoms colored white, red, blue, yellow, light orange, and orange for carbon, oxygen, nitrogen, sulfur, phosphorus, and iron, respectively.

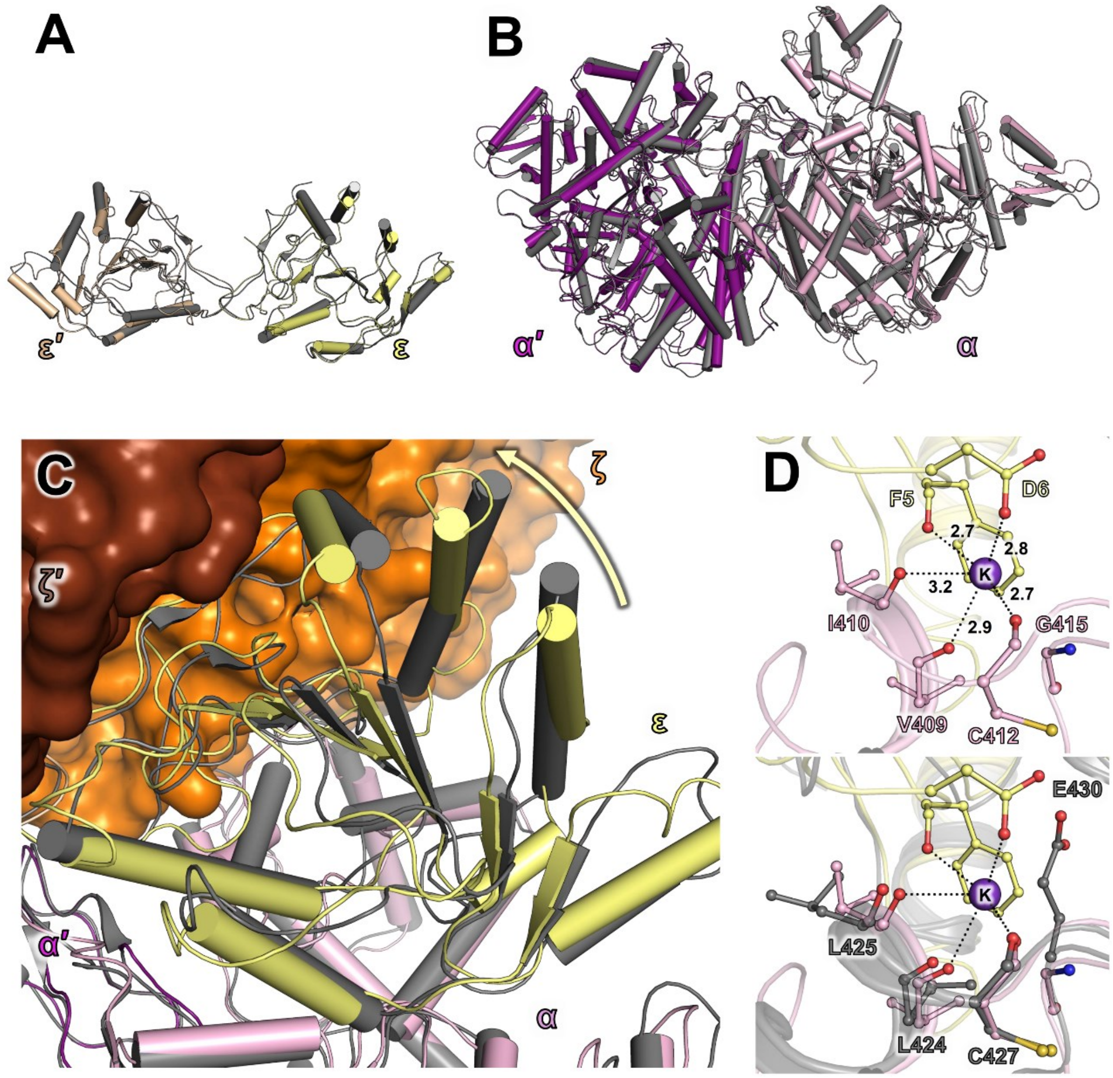

**Figure S3. Alignment of $\alpha_2\varepsilon_2\zeta_2$ subcomplex from *Ca.* E. thermophilum and the $\alpha_2\varepsilon_2$ subcomplex from *M. barkeri.*** **A-B.** Superposition of the structures of the ACDS $\alpha_2\varepsilon_2\zeta_2$ subcomplex and its equivalent from *M. barkeri* (PDB 3CF4 (27)). **A.** Superimposition of the structure of the ε subunit from the *M. barkeri* (grey cartoon) on both the ε and ε′ subunits of the subcomplex from *Ca*. E. thermophilum (light yellow and wheat cartoon, respectively). **B**. Superimposition of the structure of the αα′ core from the *M. barkeri* (grey cartoon) on the αα′ core of the subcomplex from *Ca*. E. thermophilum (light pink and deep purple, respectively). The r.m.s.d. and coverage are given in Table S3. **C.** The $\alpha_2\varepsilon_2$ subcomplex from *Ca*. E. thermophilum and *M. barkeri* are shown as cartoons, colored as in **A**, and aligned on the αα′ core. The ζζ′ dimer is shown as surface, with subunits colored in orange and brown, respectively. The movement of the ε subunit triggered by the presence of the ζ subunits is shown as a yellow arrow. **D.** Top: Coordination of the ion modelled as a K atom (purple sphere) in the structure from *Ca.* E. thermophilum at the interface of the α (pink) and ε (light yellow) subunits. Bottom: superimposition with the structure of the subcomplex from *M. barkeri* (grey). Proteins are shown as cartoons and the residues in the vicinity are shown as balls and sticks with oxygen, nitrogen and sulfur colored red, blue and yellow, respectively, with contacts shown as black dashes.

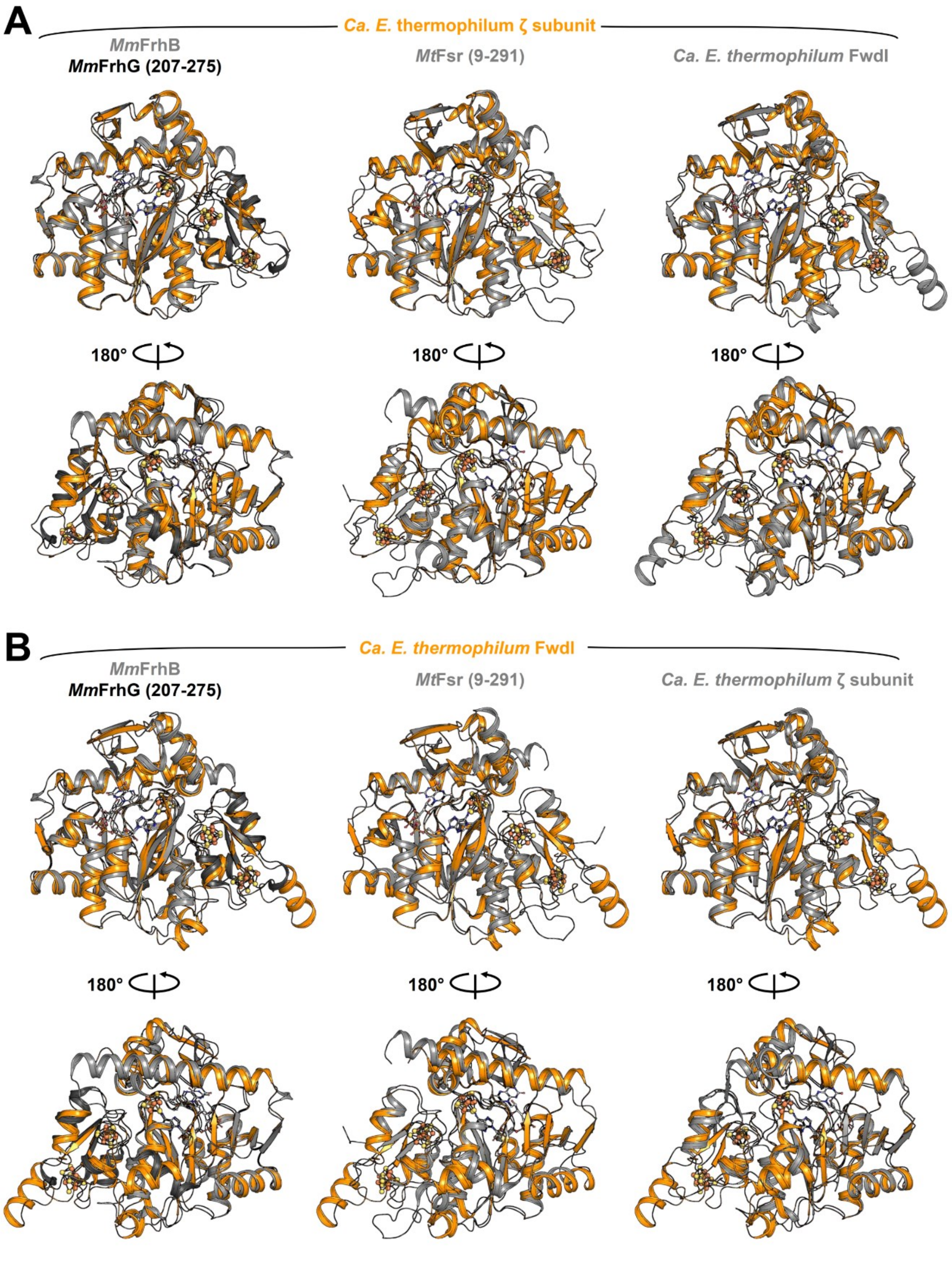

**Figure S4. Alignment of the ζ subunit from the ACDS and FwdI subunit from the Fwd complex of *Ca*. E. thermophilum with structurally characterized homologs. A-B.** Alignment of the ζ subunit from the ACDS (**A**) or the FwdI subunit (**B**) with the structures of the FrhBG (γ207-275) subcomplex from *M. marburgensis* (PDB 4OMF) and Fsr (9-291) from *M. thermolithotrophicus* (PDB 7NP8). The alignment r.m.s.d. and coverage are given in Table S3.

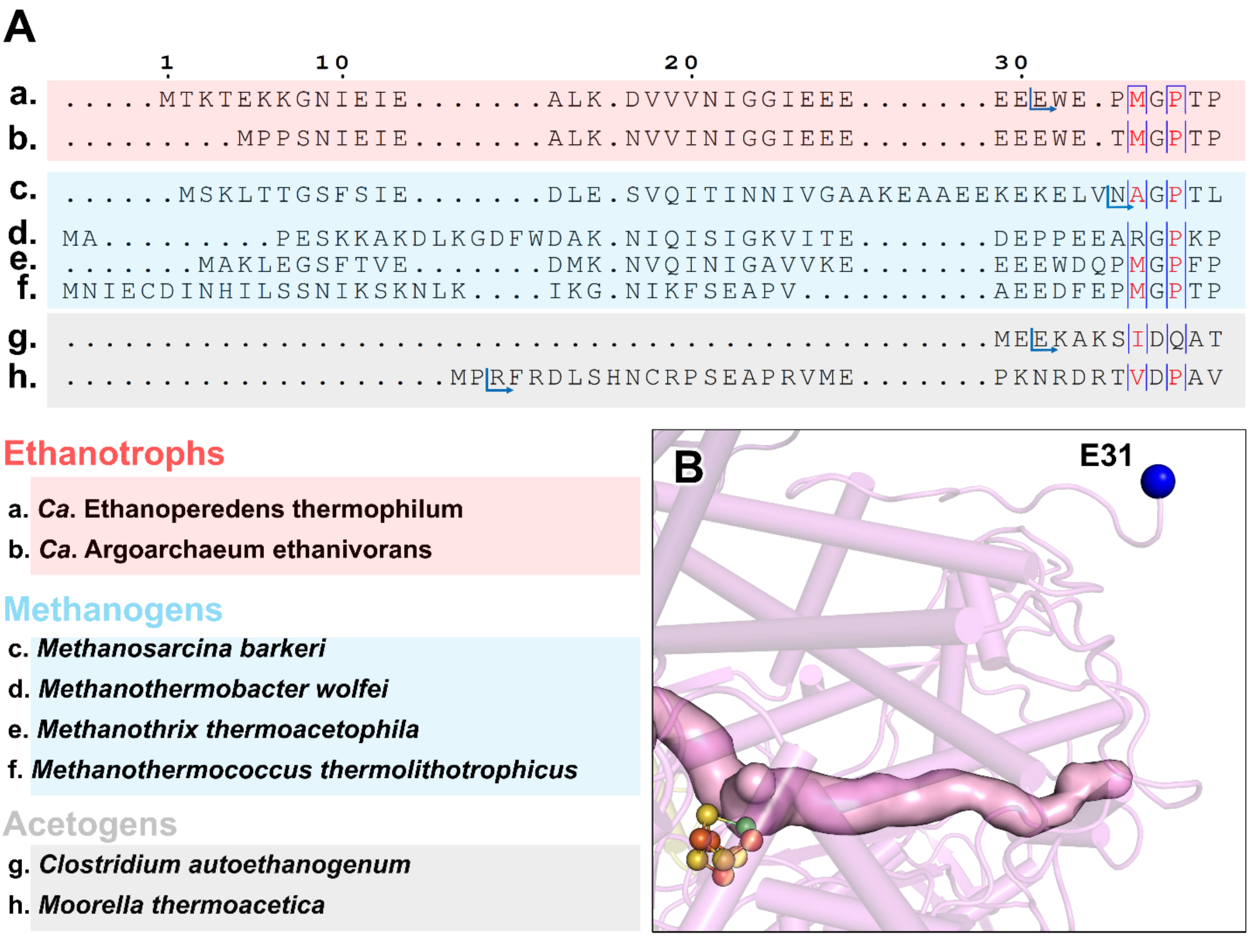

**Figure S5. N-terminal extension in CO-dehydrogenases. A.** Multiple alignments of the sequences of CODH (ACDS α-subunit) in archaea and bacteria. The numbering corresponds to the sequence of the protein from *Ca*. E. thermophilum. A blue arrow indicates the beginning of the chain in the respective structural model. **B.** Position of the first modeled residue of the α-subunit from *Ca*. E. thermophilum. The protein is shown as a cartoon, colored in light pink, with the C-cluster shown as balls and sticks with sulfur, nickel, and iron colored in yellow, green, and orange, respectively. The N-terminus of the model is represented as a blue ball. The CO-diffusion channel predicted by the CAVER program is shown as a light pink surface.

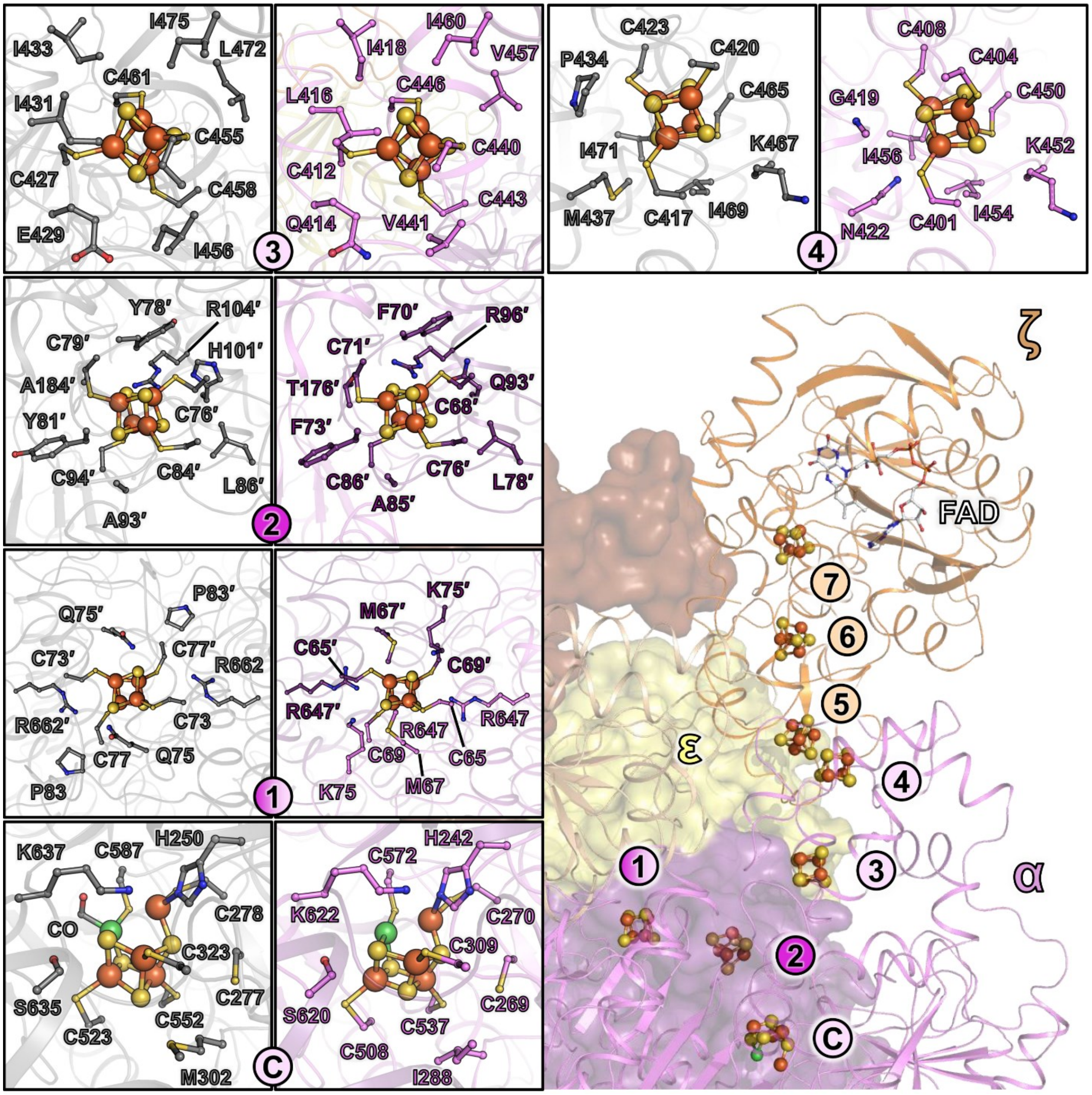

**Figure S6. Metallo-cofactors coordination in the $\alpha_2\varepsilon_2$ subcomplex from *Ca*. E. thermophilum.** The α, ε and ζ subunits are represented as cartoons and the α′, ε′ and ζ′ subunits as a surface to provide an overall view of the metallo-cofactor locations. Framed panels display a close-up view of the coordination comparing the $\alpha_2\varepsilon_2$-subcomplex from *M. barkeri* (PDB 3CF4, grey) and *Ca*. E. thermophilum (colored as in the overall view). The different cofactors and residues in their vicinity are represented in balls and stick with nitrogen, oxygen, sulfur, phosphorus, nickel, and iron colored in blue, red, yellow, light orange, green, and orange. Carbon atoms of the FAD cofactor are colored white. The coordination of the cofactors from the ζ subunit is described in Figure S8.

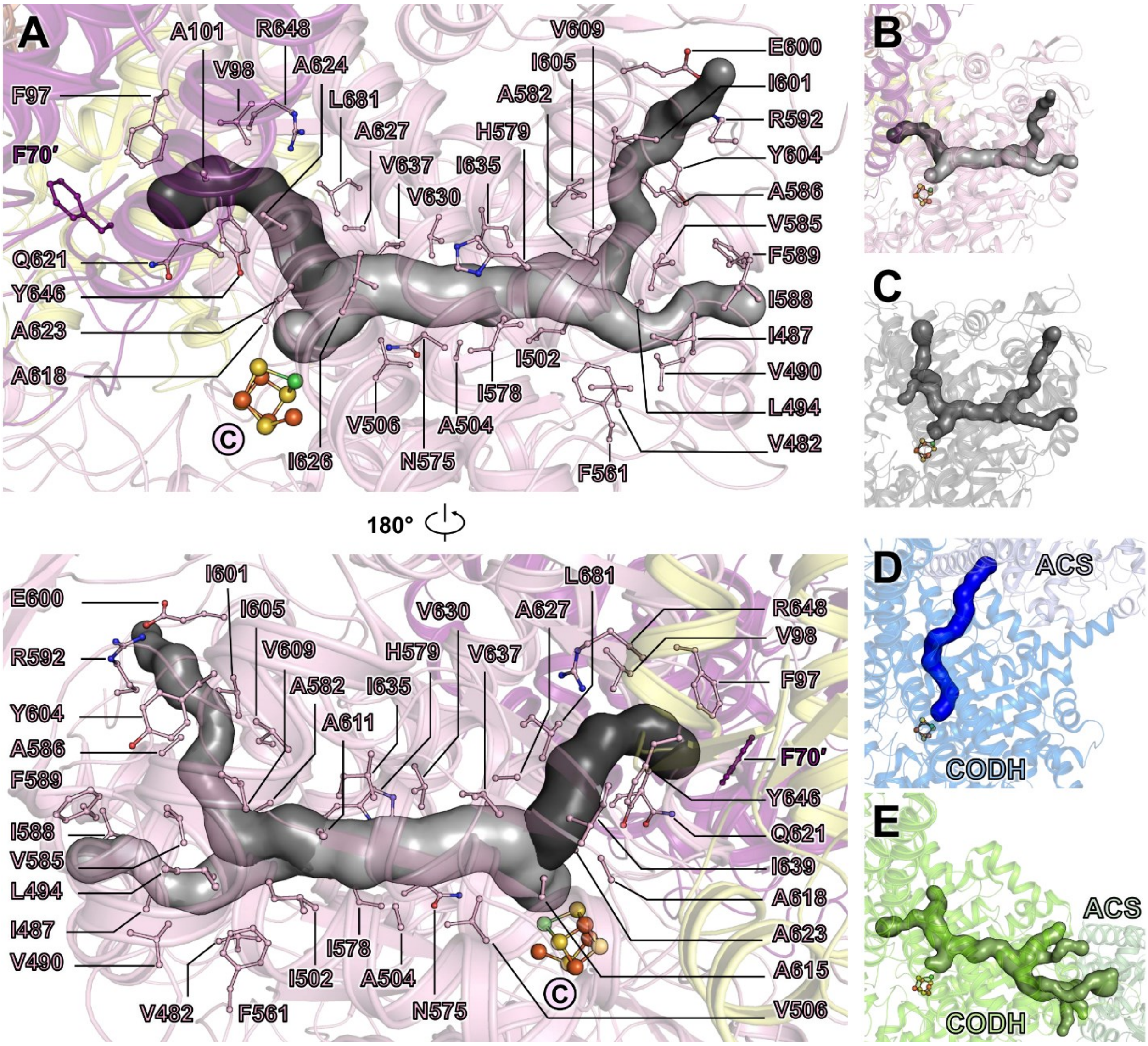

**Figure S7. Tunneling systems of the $\alpha_2\varepsilon_2\zeta_2$ ACDS from *Ca*. E. thermophilum. A.** The different tunnels predicted by the CAVER program (interrupted at the hydrophilic bottleneck near F70′), shown as surface, and colored in shades of grey. The proteins are shown in cartoons with α, α′ and ε subunits colored in light pink, deep purple, and light yellow, respectively. The residues forming the channel and cofactors are shown in balls and sticks with oxygen, nitrogen, sulfur, iron, and nickel atoms colored in red, blue, yellow, orange, and green, respectively. **B-E.** Tunneling systems comparison between the $\alpha_2\varepsilon_2\zeta_2$ ACDS from *Ca.* E. thermophilum (**B**), the $\alpha_2\varepsilon_2$ ACDS from *M. barkeri* (PDB 3CF4, **C**), the CODH/acetyl-coenzyme A (ACS) from *Moorella thermoacetica* (PDB 1MJG, **D**), and the CODH/ACS from *Clostridium autoethanogenum* (PDB 6YTT, **E**). The C-cluster is shown as balls and sticks with sulfur, iron, and nickel colored in yellow, orange, and green, respectively. **D and E.** The structures are colored in shades of blue and green, respectively, with the CODH in lighter and ACS darker colors.

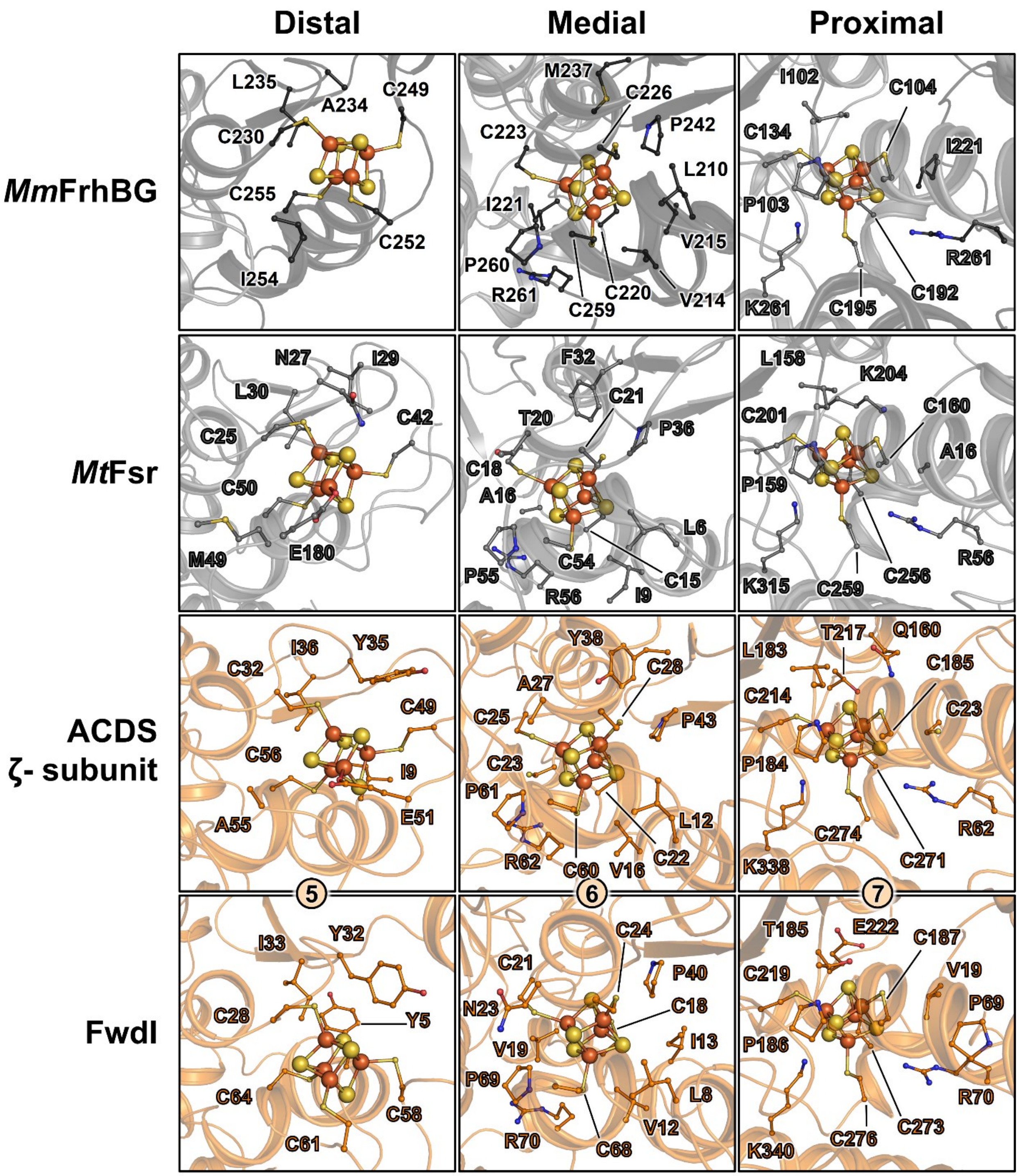

**Figure S8. Metallo-cofactor coordination in the $F_{420}$ reductases from *Ca*. E. thermophilum and related structures.** The coordination of the distal, medial and proximal clusters in the *Mm*FrhBG structure (PDB 4OMF, colored in grey and black, 1st raw), *Mt*Fsr (PDB 7NP8, colored in grey, 2nd raw), the ACDS ζ subunit (colored in orange, 3rd raw) and the FwdI subunit of the Fwd complex (colored in orange, 4th raw) is shown. The proteins are displayed as cartoons, with the clusters and the surrounding residues shown as balls and sticks with oxygen, nitrogen, sulfur, and iron colored in red, blue, yellow, and orange, respectively.

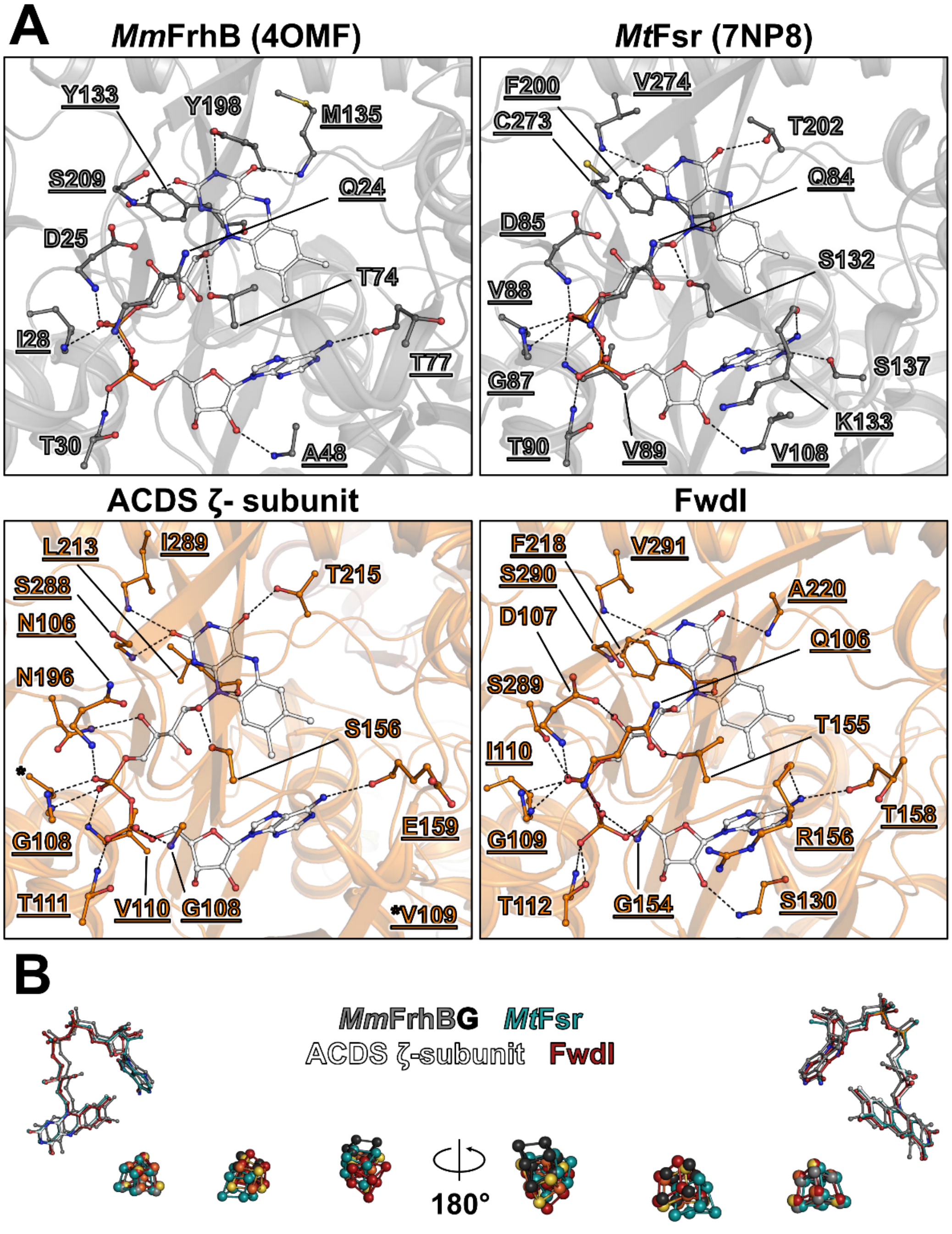

**Figure S9. FAD coordination in the F$_{420}$ reductases from *Ca.* E. thermophilum and related structures. A.** Proteins are displayed in cartoons, colored grey (*Mm*FrhB and *Mt*Fsr) or orange (ACDS ζ subunit and FwdI). FAD is colored white. The residues surrounding the FAD are shown as balls and sticks, with contacts shown as black dashes. Residues in contact with the FAD by their main or side chain are labelled underlined or not, respectively (and not underlined if both). Oxygen, nitrogen, sulfur and phosphorus are colored red, blue, yellow and light orange, respectively. The carbons from the FAD are colored white. **B.** Superimposition of the (metallo)-cofactors of the different structures, colored as indicated. The structural alignment was done using the complete ACDS ζ subunit, the complete FwdI subunit, the FrhBG (γ207-275) subcomplex (PDB 4OMF) and Fsr (9-291, PDB 7NP8). In the ACDS ζ subunit atoms are colored per type, as in A.

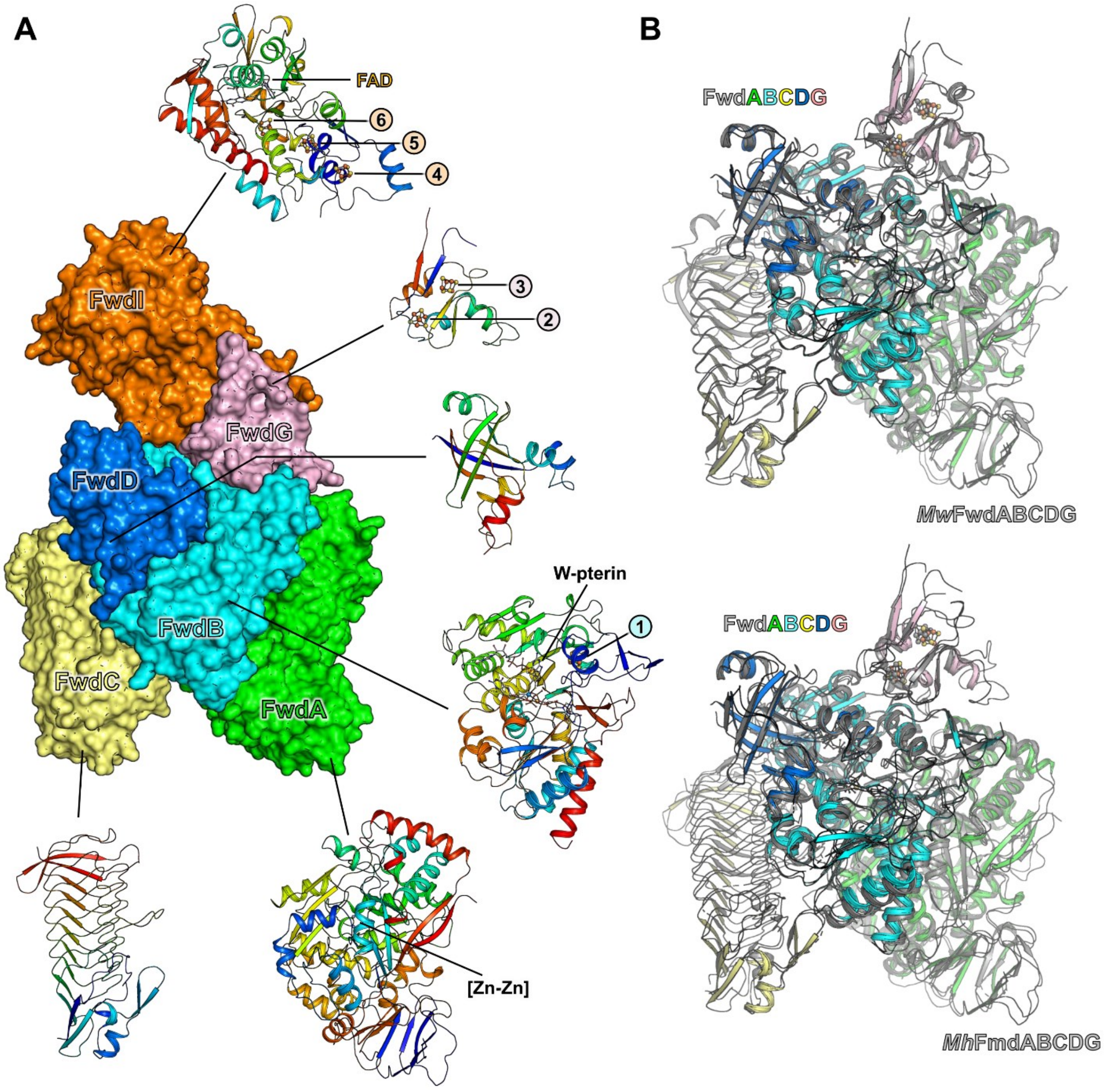

**Figure S10. Structure of the individual components of the Fwd complex from *Ca*. E. thermophilum and superposition with related structures. A.** Structure of the Fwd complex (surface) and the isolated subunits (cartoon). The isolated subunits are colored from the N-terminus to the C-terminus in a rainbow gradient from blue to red, in the same pose as the top left panel. Cofactors are shown as balls and sticks with carbon, oxygen, nitrogen, sulfur, phosphorus, iron, zinc and tungsten colored in white, red, blue, yellow, light orange, orange, grey and greyish blue, respectively. **B.** Alignment of the Fwd complex from *Ca*. E. thermophilum (colored as in **A**), the FwdABCDG core of the Fwd complex from *M. wolfei* (PDB 5T5M, colored in grey, top), and the FmdABCDG core of the Fmd-containing supercomplex from *M. hungatei* (PDB 7BKB, colored in grey, bottom). The proteins are shown as cartoons with cofactors shown as balls and sticks and colored as in **A**. The alignment r.m.s.d. and coverage are given in Table S3.

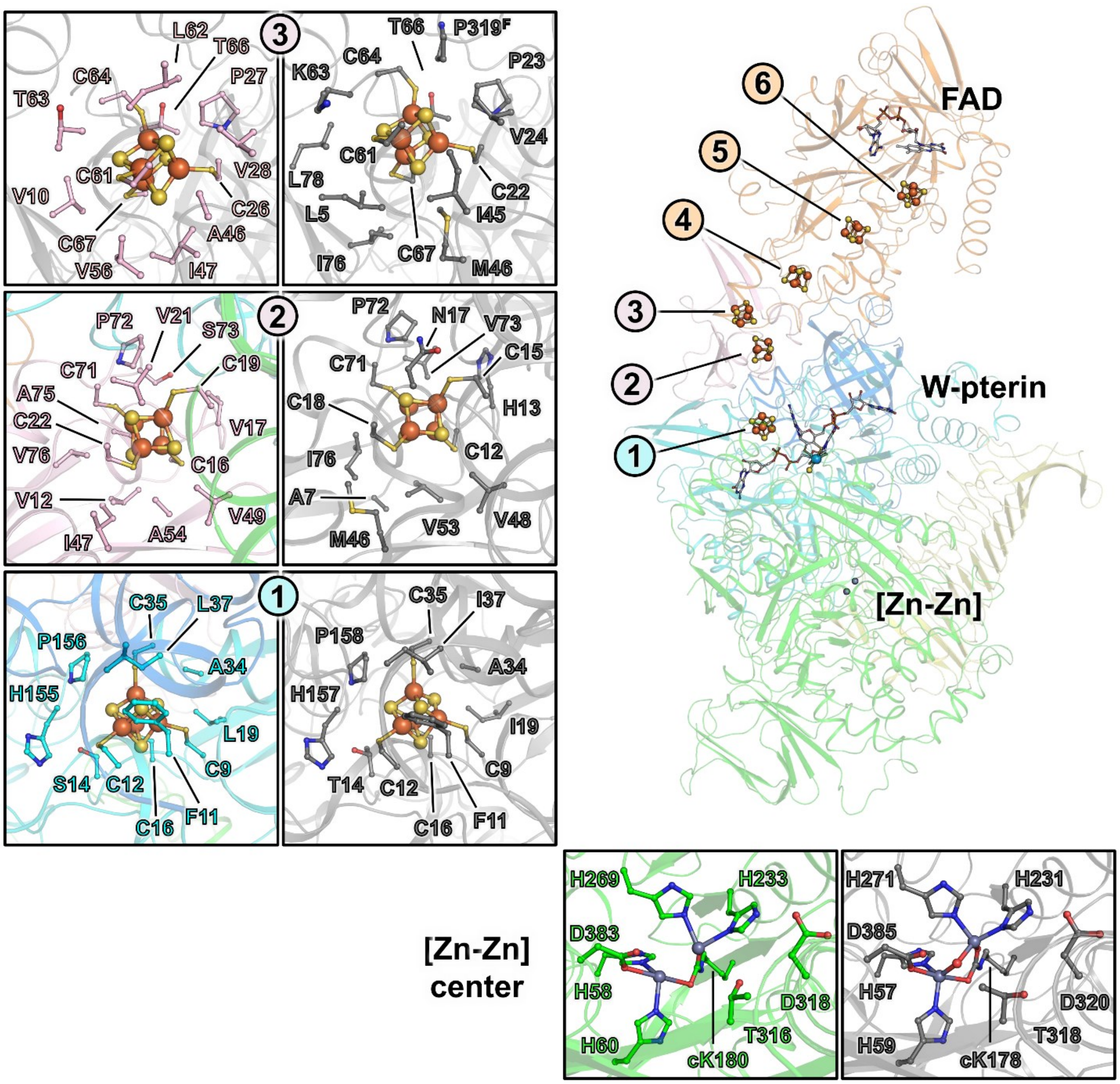

**Figure S11. Cofactors coordination in the Fwd complex from *Ca*. E. thermophilum and *M. wolfei*.** The proteins are represented as cartoons with the A, B, C, D, G, and I subunits of the Fwd complex from *Ca*. E. thermophilum colored green, cyan, light yellow, marine blue, light pink, and orange, respectively, and the proteins from *M. wolfei* colored grey. The different cofactors and residues in their vicinity are represented in balls and sticks with nitrogen, oxygen, sulfur, phosphorus, zinc, tungsten, and iron colored in blue, red, yellow, light orange, grey, grey-blue, and orange. Carbon atoms of the FAD and tungstopterin cofactors are colored white. The P319$^{F}$ is labelled "$^{F}$" because the residue is part of the FwdF subunit instead of FwdG.

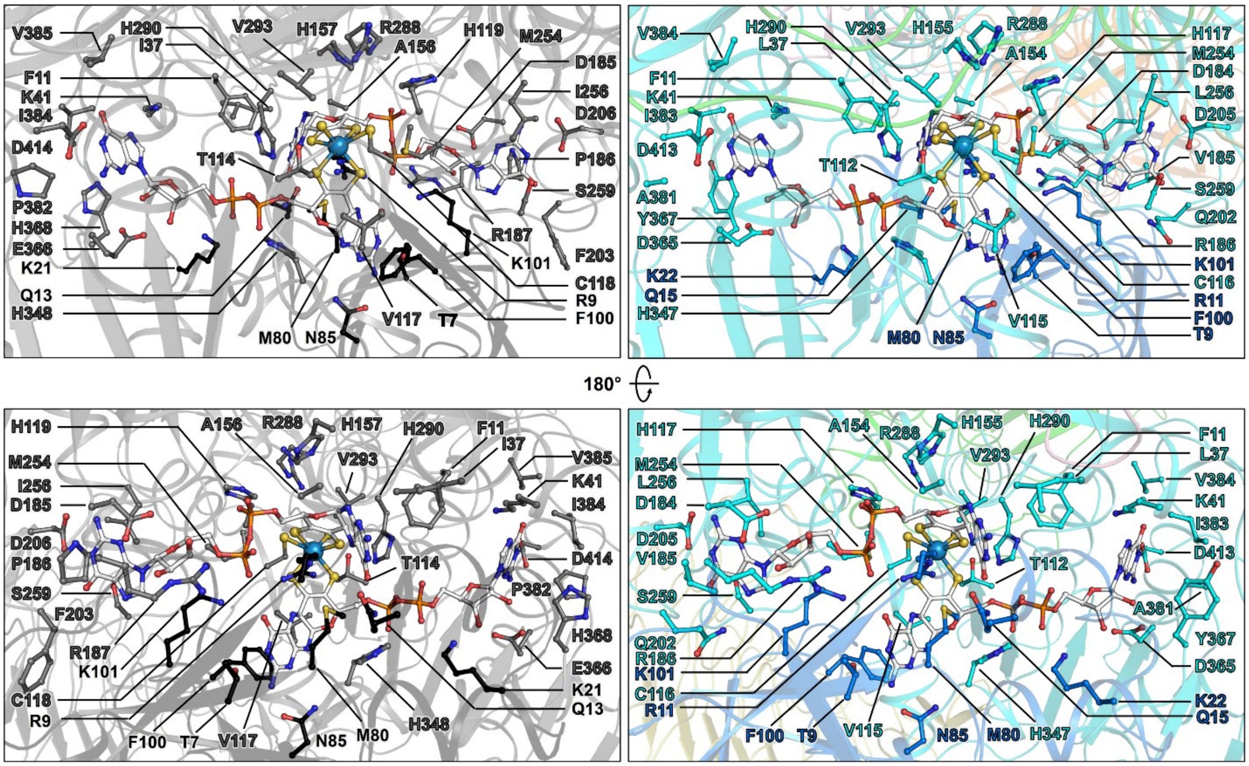

**Figure S12. Tungstopterin coordination in the Fwd complex from *Ca*. E. thermophilum and *M. wolfei*.** The proteins are shown as cartoons with the A, B, C, D, G, and I subunits of the Fwd complex from *Ca*. E. thermophilum colored in green, cyan, light yellow, marine blue, light pink, and orange, respectively, and all subunits from *M. wolfei* colored in grey except for FwdD colored black. The tungstopterin and residues in its vicinity are represented in balls and sticks with nitrogen, oxygen, sulfur, phosphorus, and tungsten colored in blue, red, yellow, light orange and grey blue. Carbon atoms of the tungstopterin are colored white.

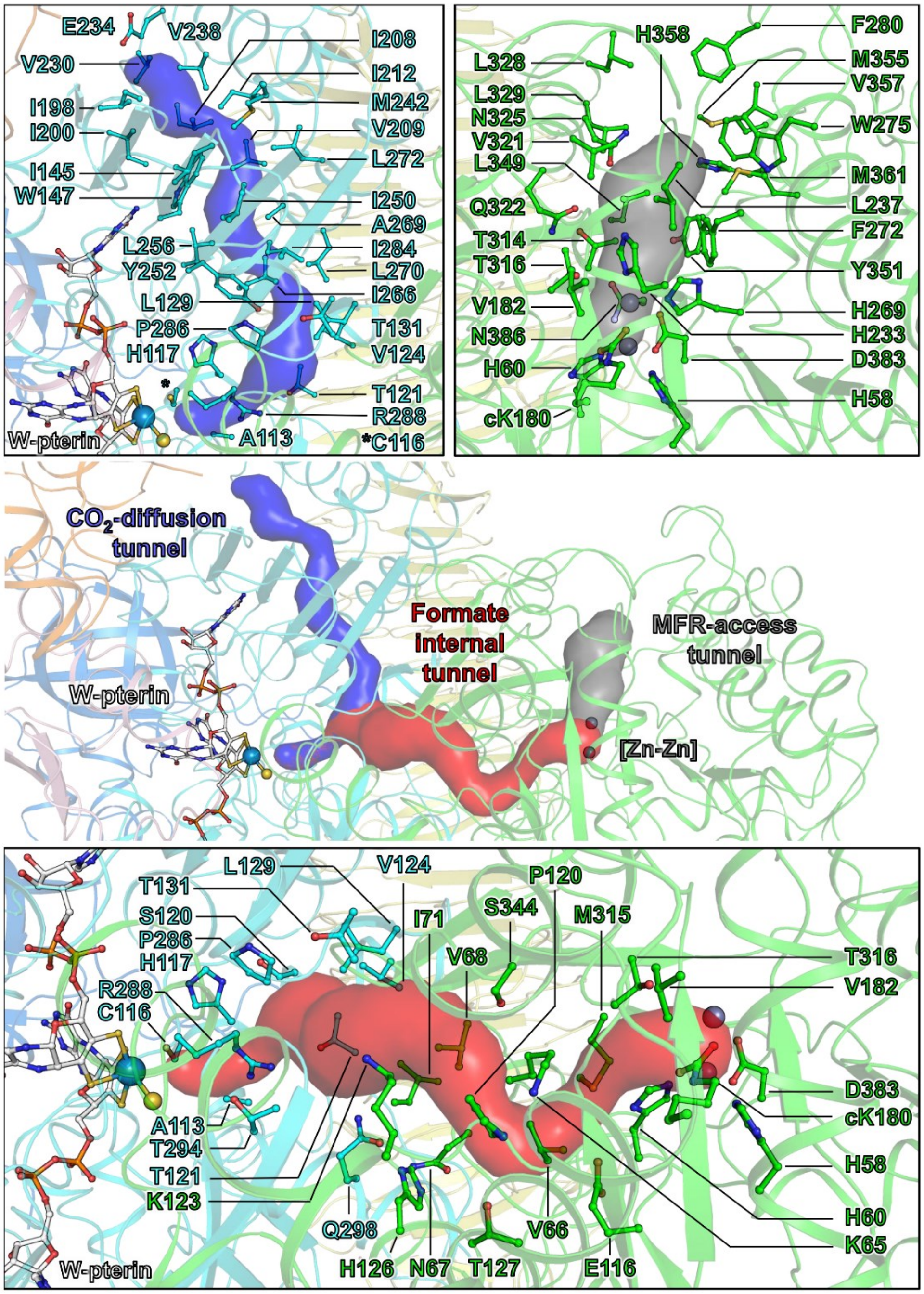

**Figure S13. Tunneling systems of the Fwd complex from *Ca*. E. thermophilum.** The central panel presents the overall structure and the different functional tunnels predicted by the CAVER program (displayed as transparent surfaces). Details of the tunnel composition are shown in the framed subpanels. The subunits are represented as cartoons with the A, B, C, D, G, and I subunits colored green, cyan, light yellow, marine blue, light pink, and orange, respectively. The residues structuring each tunnel and metallo-cofactors are displayed as balls and sticks with oxygen, nitrogen, sulfur, phosphorus, zinc, and tungsten atoms colored red, blue, yellow, light orange, dark grey, and greyish blue, respectively. The carbon atoms of the tungstopterin are colored white.

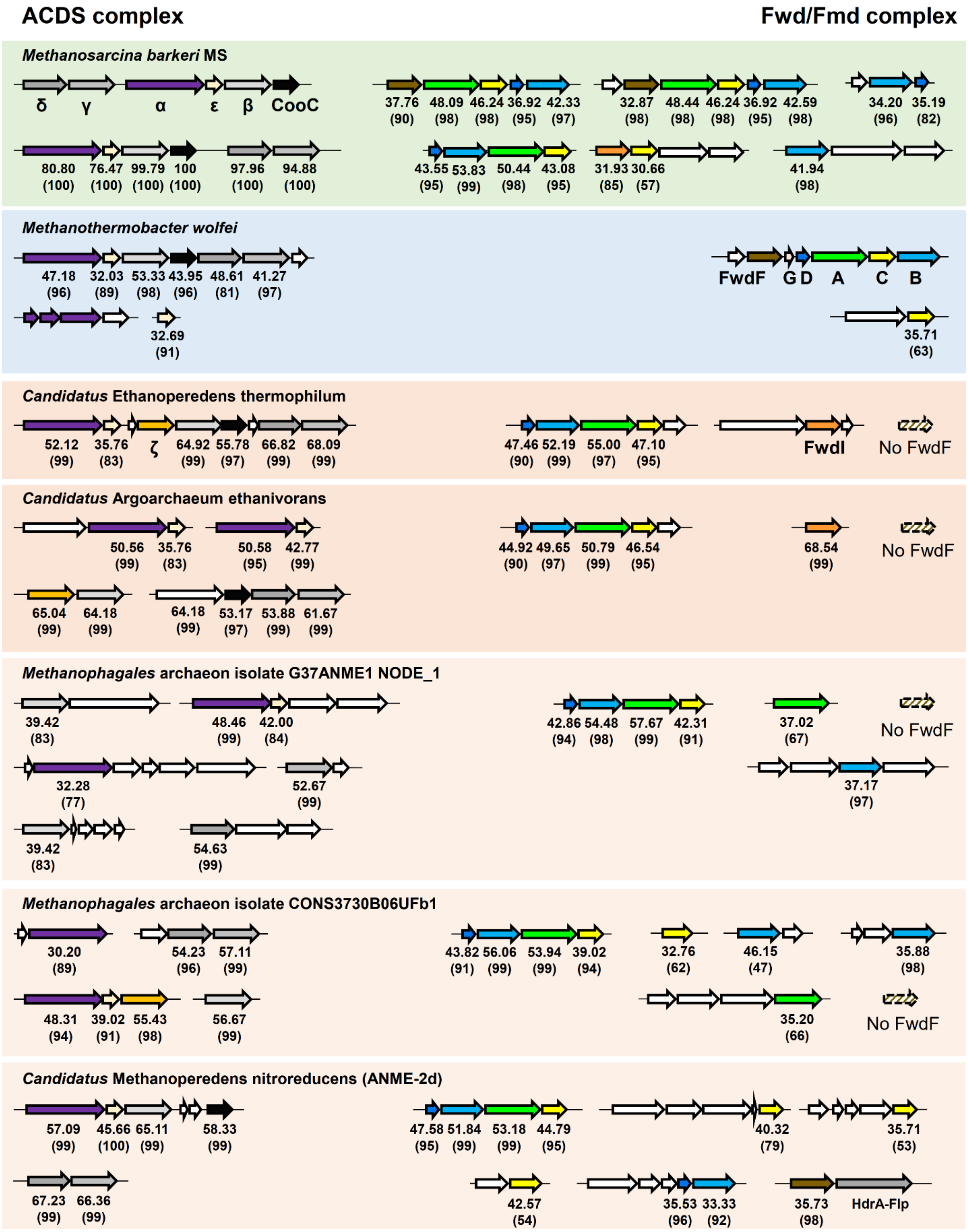
ACDS complex
Fwd/Fmd complex
Methanosarcina barkeri MS
δ γ α ε β CooC
37.76 (90) 48.09 (98) 46.24 (98) 36.92 (95) 42.33 (97)
32.87 (98) 48.44 (98) 46.24 (98) 36.92 (95) 42.59 (98)
34.20 (96) 35.19 (82)
80.80 (100) 76.47 (100) 99.79 (100) 100 (100) 97.96 (100) 94.88 (100)
43.55 (95) 53.83 (99) 50.44 (98) 43.08 (95)
31.93 (85) 30.66 (57)
41.94 (98)
Methanothermobacter wolfei
47.18 (96) 32.03 (89) 53.33 (98) 43.95 (96) 48.61 (81) 41.27 (97)
FwdF G D A C B
32.69 (91)
35.71 (63)
Candidatus Ethanoperedens thermophilum
52.12 (99) 35.76 (83) ζ 64.92 (99) 55.78 (97) 66.82 (99) 68.09 (99)
47.46 (90) 52.19 (99) 55.00 (97) 47.10 (95)
FwdI
No FwdF
Candidatus Argoarchaeum ethanivorans
50.56 (99) 35.76 (83) 50.58 (95) 42.77 (99)
44.92 (90) 49.65 (97) 50.79 (99) 46.54 (95)
68.54 (99)
No FwdF
65.04 (99) 64.18 (99) 64.18 (99) 53.17 (97) 53.88 (99) 61.67 (99)
Methanophagales archaeon isolate G37ANME1 NODE_1
39.42 (83) 48.46 (99) 42.00 (84)
42.86 (94) 54.48 (98) 57.67 (99) 42.31 (91)
37.02 (67)
No FwdF
32.28 (77) 52.67 (99)
37.17 (97)
39.42 (83) 54.63 (99)
Methanophagales archaeon isolate CONS3730B06UFb1
30.20 (89) 54.23 (96) 57.11 (99)
43.82 (91) 56.06 (99) 53.94 (99) 39.02 (94)
32.76 (62)
46.15 (47)
35.88 (98)
48.31 (94) 39.02 (91) 55.43 (98) 56.67 (99)
35.20 (66)
No FwdF
Candidatus Methanoperedens nitroreducens (ANME-2d)
57.09 (99) 45.66 (100) 65.11 (99) 58.33 (99)
47.58 (95) 51.84 (99) 53.18 (99) 44.79 (95)
40.32 (79)
35.71 (53)
67.23 (99) 66.36 (99)
42.57 (54)
35.53 (96) 33.33 (92)
35.73 (98) HdrA-Flp

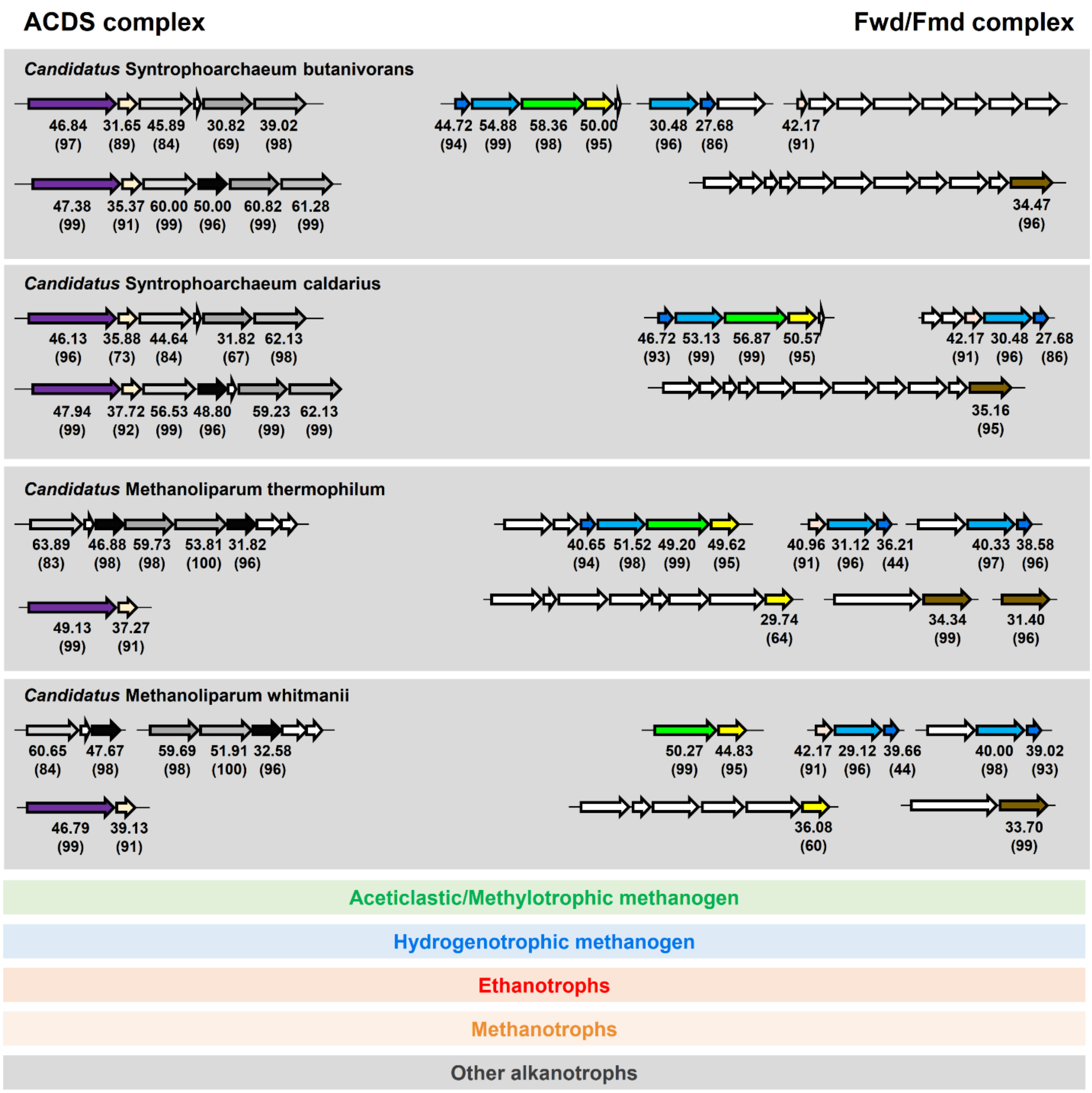

**Figure S14. Genomic environment of genes coding for ACDS and Fmd/Fwd subunits in methanogens, methanotrophs, ethanotrophs, and other alkanotrophs.** Genes are represented by arrows with size depending on the gene length. The genes coding for the α, ε, β, γ, δ, and ζ subunits of ACDS colored deep purple, light yellow, light grey, grey, dark grey, and orange, respectively, the maturation protein CooC in black, and the A, B, C, D, G, F and I subunits of the Fmd/Fwd complex in green, cyan, light yellow, marine blue, light pink, brown and dark orange, respectively. Other genes are colored white. Gene annotation is based on sequence identity as determined by BLAST. The percentage of identity (and percentage of coverage) are given. Putative operon organization, suggested by the Operon Mapper webserver (26), is represented by a continuous line. The background is colored according to the main metabolism of the organism. The analysis presents a putative $F_{420}$-reducing CODH in some ANME-1 species because of the presence of genes coding for a putative protein similar to the ζ subunit from the ethanotrophs.

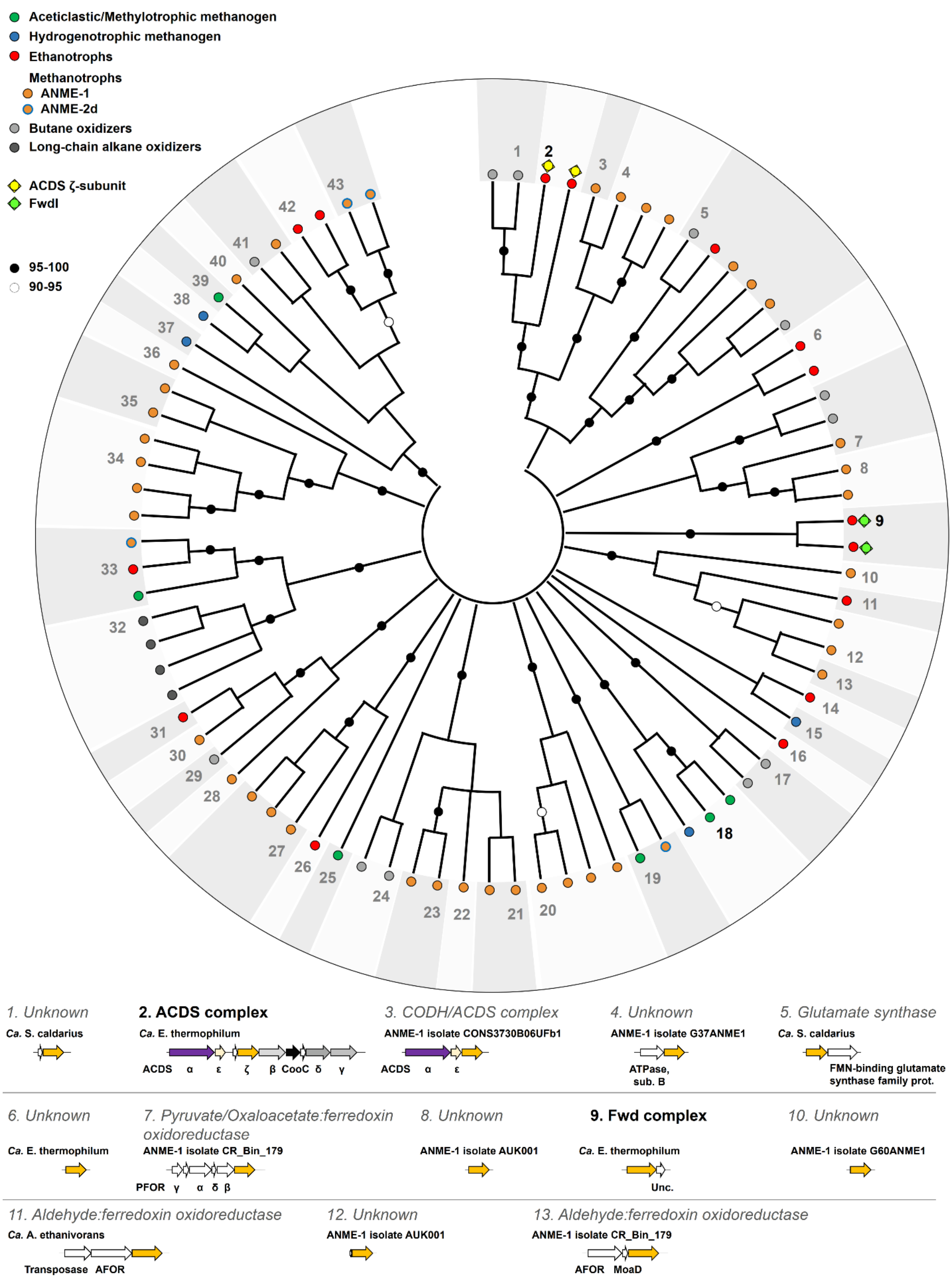
Aceticlastic/Methylotrophic methanogen
Hydrogenotrophic methanogen
Ethanotrophs
Methanotrophs
ANME-1
ANME-2d
Butane oxidizers
Long-chain alkane oxidizers
ACDS ζ-subunit
FwdI
95-100
90-95
1. Unknown
Ca. S. caldarius
2. ACDS complex
Ca. E. thermophilum
ACDS α ε ζ β CooC δ γ
3. CODH/ACDS complex
ANME-1 isolate CONS3730B06UFb1
ACDS α ε
4. Unknown
ANME-1 isolate G37ANME1
ATPase, sub. B
5. Glutamate synthase
Ca. S. caldarius
FMN-binding glutamate synthase family prot.
6. Unknown
Ca. E. thermophilum
7. Pyruvate/Oxaloacetate:ferredoxin oxidoreductase
ANME-1 isolate CR_Bin_179
PFOR γ α δ β
8. Unknown
ANME-1 isolate AUK001
9. Fwd complex
Ca. E. thermophilum
Unc.
10. Unknown
ANME-1 isolate G60ANME1
11. Aldehyde:ferredoxin oxidoreductase
Ca. A. ethanivorans
Transposase AFOR
12. Unknown
ANME-1 isolate AUK001
13. Aldehyde:ferredoxin oxidoreductase
ANME-1 isolate CR_Bin_179
AFOR MoaD

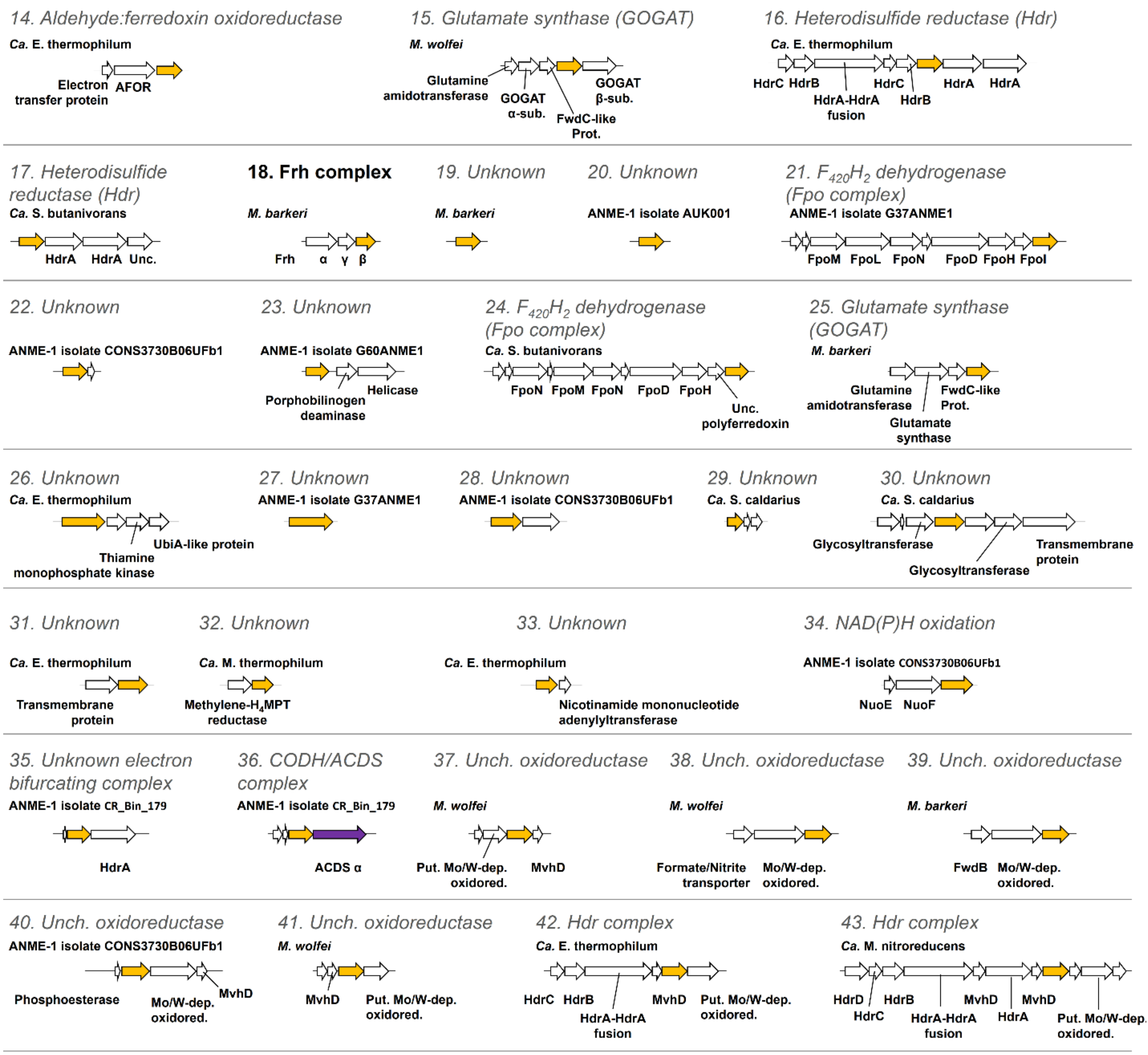

**Figure S15. Phylogenic tree of F420 reductase homologs and respective gene environment in a selection of methanogens, methanotrophs, ethanotrophs, and other alkanotrophs.** All sequences of $F_{420}H_2$ oxidases homologous to the ζ/FwdI subunits extracted from the genome of various organisms were used to construct a Maximum-likelihood evolution phylogenic tree (top panel). Subgroups of the tree were established according to the genomic environment, shown in the bottom panel. Genes are represented by arrows with size depending on the gene length and gene coding for the α, ε, β, γ, and δ subunits of ACDS colored deep purple, light yellow, light grey, grey, dark grey, and orange, respectively, the maturation protein CooC in black and homologs of the and ζ subunit of ACDS in orange. Other genes are colored white. Gene annotation is based on sequence identity determined by BLAST using the SwissProt and PDB databases. Putative operon organization, suggested by the Operon Mapper webserver (26), is represented by a continuous line. “prot.” “sub.”, “unc.”, “put.” and “oxidored.” stand for protein, subunit, uncharacterized, putative and oxidoreductase, respectively. The ζ subunit homolog encoded in the genome of *Ca.* Syntrophoarchaeum species may suggest a $F_{420}$-reducing ACDS for one of the two ACDS isoforms of the organisms (Fig. S14).

**Table S1. Purification of the CODH component and Fwd of *Ca*. E. thermophilum.**

| Fraction, purification step | Activity (µmol of CO oxidized per minute) | Protein (mg) | Specific activity (µmol of CO oxidized per minute per mg of protein) | Yield (%) | Purification (fold) |
|---|---|---|---|---|---|
| **CODH component** | | | | | |
| **Soluble extract** | 90.83 ± 7.40 | 105.42 | 0.86 ± 0.07 | 100 | 1 |
| **Q-Sepharose** | 56.05 ± 6.38 | 48.28 | 1.16 ± 0.13 | 61.71 | 1.35 |
| **Source 15PHE** | 21.92 ± 1.63 | 2.78 | 7.88 ± 0.59 | 24.13 | 9.15 |
| **Source 15PHE** | 19.24 ± 2. 78 | 2.07 | 9.30 ± 1.34 | 21.19 | 10.80 |
| **Superdex 200 10/300** | 10.50 ± 1. 51 | 0.82 | 12.80 ± 1.84 | 11.56 | 14.86 |
| **Fwd complex** | | | | | |
| **Furfurylformamide dehydrogenase activity** | | | | | |
| **Soluble extract** | 37.54 ± 7.61 | 105.42 | 0.36 ± 0.07 | 100 | 1 |
| **Q-Sepharose** | 5.90 ± 0.64 | 48.28 | 0.12 ± 0.01 | 15.73 | 0.34 |
| **Source 15PHE** | 2.47 ± 0.12 | 0.55 | 4.46 ± 0.21 | 6.57 | 12.50 |
| **Superdex 200 10/300** | 2.02 ± 0.11 | 0.40 | 5.05 ± 0.27 | 5.38 | 14.18 |
| **Formate dehydrogenase activity** | | | | | |
| **Soluble extract** | 130.27 ± 10.27 | 105.42 | 1.24 ± 0.10 | 100 | 1 |
| **Q-Sepharose** | 3.59 ± 0.33 | 48.28 | 0.07 ± 0.01 | 2.76 | 0.06 |
| **Source 15PHE** | 0.07 ± 0.01 | 0.55 | 0.12 ± 0.01 | 0.05 | 0.10 |
| **Superdex 200 10/300** | 0.05 ± 0.01 | 0.40 | 0.17 ± 0.06 | 0.04 | 0.14 |

**Table S2. X-ray analysis statistics for the structures obtained from *Ca*. E. thermophilum.**

| | $\alpha_2\varepsilon_2\zeta_2$ ACDS subcomplex SAD Fe K-edge | $\alpha_2\varepsilon_2\zeta_2$ ACDS subcomplex | Fwd complex |
|---|---|---|---|
| **Data collection** | | | |
| Synchrotron source | SLS, X06DA | SOLEIL, PROXIMA-1 | SLS, X06DA |
| Wavelength (Å) | 1.73981 | 0.97856 | 1.00000 |
| Space group | $C222_1$ | $P2_12_12_1$ | $P2_1$ |
| Resolution (Å) | 57.27 −3.00 (3.05 −3.00) | 122.41 −1.89 (2.10 −1.89) | 57.38 −1.97 (2.19 −1.97) |
| Cell dimensions | | | |
| a, b, c (Å) | 109.40, 196.33, 494.09 | 97.07, 159.21, 191.44 | 107.63, 135.64, 149.90 |
| α, β, γ (°) | 90, 90, 90 | 90, 90, 90 | 90, 90.49, 90 |
| $R_{merge}$(%)[a] | 35.4 (171.5) | 14.9 (158.3) | 25.1 (120.0) |
| $R_{pim}$ (%)[a] | 9.9 (49.8) | 4.2 (46.0) | 10.1 (48.6) |
| $CC_{1/2}$ [a] | 0.987 (0.590) | 0.999 (0.639) | 0.989 (0.621) |
| $I/\sigma_I$ [a] | 7.3 (1.6) | 10.8 (1.7) | 7.2 (1.6) |
| Spherical completeness[a] | 100.0 (100.0) | 70.0 (13.0) | 55.7 (10.5) |
| Ellipsoidal completeness[a] | - | 95.9 (73.6) | 93.0 (64.3) |
| Redundancy[a] | 13.6 (12.7) | 13.6 (12.2) | 7.0 (6.9) |
| Nr. unique reflections[a] | 106,477 (5,263) | 164,775 (8,240) | 167,948 (8,397) |
| **Refinement** | | | |
| Resolution (Å) | - | 39.80 −1.89 | 57.38 −1.97 |
| Number of reflections | - | 164,728 | 167,913 |
| $R_{work}/R_{free}$[b] (%) | - | 16.51/18.73 | 17.57/21.00 |
| Number of atoms | | | |
| Protein | - | 20,153 | 27,665 |
| Solvent and ligands | - | 371 | 581 |
| Water | - | 1,418 | 2,282 |
| Mean B-value ($Å^2$) | - | 44.12 | 25.29 |
| Molprobity clash score, all atoms | - | 2.44 | 1.37 |
| Ramachandran plot | | | |
| Favored regions (%) | - | 97.88 | 97.00 |
| Outlier regions (%) | - | 0.08 | 0.11 |
| r.m.s.d.[c] bond lengths (Å) | - | 0.009 | 0.010 |
| r.m.s.d.[c] bond angles (°) | - | 1.233 | 1.339 |
| **PDB ID code** | | **8RIU** | **8RJA** |

[a] Values relative to the highest resolution shell are within parentheses. [b] $R_{free}$ was calculated as the $R_{work}$ for 5 % of the reflections that were not included in the refinement. [c] r.m.s.d., root mean square deviation.

**Table S3. Structural alignment of the $\alpha_2\varepsilon_2$ core and ζ subunit of ACDS from *Ca*. E. thermophilum and related structures.**

| Aligned structures (name, organism, PDB code, chains and residues) | Reference structure (name, organism, PDB code, chains and residues) | r.m.s.d. (Å) | Aligned Cα |
|---|---|---|---|
| α subunit ACDS, *M. barkeri*, 3CF4, A44-A803 | $\alpha_2$ ACDS subunit, *Ca*. E. thermophilum, 8RIU, A37-A787 | 0.720 | 681 |
| ε subunit ACDS, *M. barkeri*, 3CF4, B15-B167 | $\varepsilon_2$ ACDS subunit, *Ca*. E. thermophilum, 8RIU, C12-C171 | 0.949 | 119 |
| αε ACDS, *M. barkeri*, 3CF4, A44-A804 and B15-B154 | $\alpha_2\varepsilon_2$ subunits, *Ca*. E. thermophilum, 8RIU, A37-A788 and C12-C155 | 0.789 | 748 |
| FrhBG, *M. marburgensis*, 4MOF, A213-A273 and C2-C269 | ζ subunit, *Ca*. E. thermophilum, 8RIU, E16 -E74 and E84-E346 | 0.906 | 263 |
| Fsr, *M. thermolithotrophicus*, 7NP8, A9-A291 | ζ subunit, *Ca*. E. thermophilum, 8RIU, E16-E306 | 1.264 | 211 |
| FwdI, *Ca*. E. thermophilum 8RJA, E7-E349 | ζ subunit, *Ca*. E. thermophilum, 8RIU, E7-E347 | 1.056 | 228 |
| FrhBG, *M. marburgensis*, 4MOF, A209-A271 and C6-C272 | FwdI, *Ca*. E. thermophilum, 8RJA, E6-E84 and E88-E351 | 1.311 | 251 |
| Fsr, *M. thermolithotrophicus*, 7NP8, A8-A326 | FwdI, *Ca*. E. thermophilum 8RJA, E11-E351 | 1.117 | 205 |
| FwdABCDG core, *M. wolfei*, 5T5M, A3-A566; B4-B429; C4-C261 and D1-D124 | FwdABCDG core, *Ca*. E. thermophilum, 8RJA, A3-A564; B4-B428; C4-C252 and J3-J124 | 0.791 | 1223 |
| FmdABCDG core, *M. hungatei*, 7BKB, I3-I568 and L10-L436 | FwdABCDG core, *Ca*. E. thermophilum, 8RJA, A3-A564 and B9-B426 | 0.991 | 829 |

**Table S4. Expression level of genes proposed to be involved in ethanotrophy and putatively involved in $F_{420}$ reduction/$F_{420}H_2$ oxidation.** The transcriptomics data are extracted from a previous work (1). The gene coding for the ACDS ζ subunit and FwdI are in bold.

| Locus tag | Name | Gene expression (RPKM) | | Rank |
|---|---|---|---|---|
| | | Average | Standard deviation | |
| **Ethyl-Coenzyme M reductase (ECR)** | | | | |
| FHEFKHOI_01410 | *mcrB* | 7,323.18 | 640.3673 | 28 |
| FHEFKHOI_01411 | *mcrG* | 6,676.097 | 845.1444 | 18 |
| FHEFKHOI_01412 | *mcrA* | 11,354.85 | 1371.936 | 22 |
| **Acetyl-CoA decarbonylase/synthase (ACDS)** | | | | |
| FHEFKHOI_01146 | *cdhA* | 2,182.283 | 33.01558 | 77 |
| FHEFKHOI_01147 | *cdhE* | 2,917.7 | 272.9681 | 52 |
| **FHEFKHOI_01149** | ***cdhZ*** | **2,785.547** | **170.5093** | **54** |
| FHEFKHOI_01150 | *cdhB* | 2,619.38 | 239.1818 | 58 |
| FHEFKHOI_01153 | *cdhD_1* | 4,056.627 | 193.9212 | 36 |
| FHEFKHOI_01154 | *cdhG* | 6,749.933 | 660.5352 | 21 |
| **Methylenetetrahydromethanopterin reductase (Mer)** | | | | |
| FHEFKHOI_00600 | *mer* | 3,442.87 | 323.0438 | 45 |
| **Methylenetetrahydromethanopterin dehydrogenase (Mtd)** | | | | |
| FHEFKHOI_01914 | *mtd* | 2,144.567 | 187.2971 | 79 |
| **Methenyltetrahydromethanopterin cyclohydrolase (Mch)** | | | | |
| FHEFKHOI_00609 | *mch* | 2,499.277 | 224.7675 | 64 |
| **Formylmethanofuran--tetrahydromethanopterin formyltransferase (Ftr)** | | | | |
| FHEFKHOI_02106 | *ftr* | 1,874.323 | 117.5848 | 94 |
| **W-dependent Formylmethanofuran dehydrogenase (Fwd)** | | | | |
| FHEFKHOI_00470 | *fwdD* | 1,584.31 | 187.9854 | 126 |
| FHEFKHOI_00471 | *fwdB* | 956.78 | 69.29146 | 223 |
| FHEFKHOI_00472 | *fwdA* | 1,297.407 | 60.08871 | 161 |
| FHEFKHOI_00473 | *fwdC* | 1,532.313 | 123.3891 | 131 |
| **FHEFKHOI_01727** | ***fwdI*** | **3,569.98** | **424.5134** | **41** |
| FHEFKHOI_00695 | *fwdG* | 920.5967 | 127.3583 | 236 |
| **$F_{420}H_2$:quinone oxidoreductase (Fpo)** | | | | |
| FHEFKHOI_00303 | *fpoA* | 1,591.543 | 221.9746 | 125 |
| FHEFKHOI_00304 | *fpoB* | 3,863.963 | 496.85 | 39 |
| FHEFKHOI_00305 | *fqoC* | 3,284.12 | 377.0664 | 47 |
| FHEFKHOI_00306 | *fqoD* | 1,486.367 | 103.957 | 138 |
| FHEFKHOI_00307 | *fpoH* | 1,684.537 | 52.04053 | 113 |
| FHEFKHOI_00308 | *fpoI* | 1,482.577 | 123.4509 | 139 |
| FHEFKHOI_00309 | *fpoJ* | 1,599.91 | 268.1379 | 124 |
| FHEFKHOI_00310 | CDS | 1,883.593 | 172.2733 | 93 |
| FHEFKHOI_00311 | *fpoK* | 2,104.413 | 179.1633 | 82 |

| | | | | |
|---|---|---|---|---|
| FHEFKHOI_00312 | *fpoL_1* | 1,531.387 | 62.85304 | 132 |
| FHEFKHOI_00313 | *fpoM_1* | 1,864.007 | 59.88921 | 97 |
| FHEFKHOI_00314 | *fpoN* | 2,646.033 | 68.66652 | 57 |
| FHEFKHOI_00315 | *fpoO* | 477.6033 | 87.51585 | 486 |
| **Electron-bifurcating heterodisulfide reductase (Hdr)** | | | | |
| FHEFKHOI_01206 | *hdrA_2* | 1,252.033 | 35.98648 | 164 |
| FHEFKHOI_01207 | *hdrA_3* | 1,626.09 | 228.1921 | 117 |
| FHEFKHOI_01208 | *frhB* | 1,181.287 | 58.25349 | 181 |
| FHEFKHOI_01209 | *hdrB_2* | 1,281.423 | 96.00225 | 162 |
| FHEFKHOI_01210 | *hdrC_2* | 1,267.837 | 147.109 | 163 |
| FHEFKHOI_01211 | *hdrA_4* | 1,370.01 | 78.02363 | 152 |
| FHEFKHOI_01212 | *hdrB_3* | 853.58 | 70.98938 | 261 |
| FHEFKHOI_01213 | *hdrC_3* | 1,180.747 | 174.8528 | 182 |
| **Aldehyde oxidoreductase (Aor)** | | | | |
| FHEFKHOI_02073 | *dmsB* | 501.2867 | 85.31422 | 465 |
| FHEFKHOI_02074 | *aor_3* | 1,623.69 | 91.87875 | 118 |
| FHEFKHOI_02075 | *frhB* | 491.7167 | 76.1402 | 472 |
| **Unknown** | | | | |
| FHEFKHOI_01017 | *fpoF* | 1,449.643 | 159.1757 | 145 |
| FHEFKHOI_01018 | CDS | 651.22 | 87.53336 | 347 |
| **Glutamate synthase Glutamine oxoglutarate aminotransferase (GOGAT)** | | | | |
| FHEFKHOI_01566 | CDS | 361.7333 | 45.57154 | 627 |
| FHEFKHOI_01567 | CDS | 292.99 | 34.71367 | 754 |
| FHEFKHOI_01568 | *frhB* | 181.48 | 19.39031 | 1029 |
| **Unknown** | | | | |
| FHEFKHOI_01073 | *frhB* | 299.22 | 21.00763 | 742 |
| **Unknown** | | | | |
| FHEFKHOI_00063 | *frhB* | 483.47 | 24.79062 | 482 |
| FHEFKHOI_00064 | *mqnD* | 415.7167 | 16.60141 | 549 |
| FHEFKHOI_00065 | *thiL_1* | 321.4333 | 34.62789 | 691 |
| FHEFKHOI_00066 | *ubiA_1* | 31.58 | 6.688101 | 1899 |
| **Unknown** | | | | |
| FHEFKHOI_01857 | CDS | 23.35667 | 4.415001 | 1967 |
| FHEFKHOI_01858 | *frhB* | 68.09333 | 6.015765 | 1623 |
